# RAS-induced transformation of mammary epithelial cells relies on ZEB1-dependent cellular reprogramming via a paracrine process

**DOI:** 10.1101/2022.06.23.497180

**Authors:** Hadrien De Blander, Laurie Tonon, Frédérique Fauvet, Roxane M. Pommier, Christelle Lamblot, Rahma Benhassoun, Francesca Angileri, Benjamin Gibert, Maria Ouzounova, Anne-Pierre Morel, Alain Puisieux

## Abstract

While uncommon in breast cancers, oncogenic activation of the RAS/MAPK signalling pathway is frequent in claudin-low (CL) tumours, a subtype of breast malignancies enriched in features of epithelial-mesenchymal transition (EMT), suggesting an interplay between RAS activation and EMT. Using inducible models of human mammary epithelial cells, we show that RAS-mediated transformation relies on cellular reprogramming governed by the EMT-inducing transcription factor ZEB1. The path to ZEB1 induction involves a paracrine process: cells entering a senescent state following RAS activation release proinflammatory cytokines, notably IL-6 and IL-1α which promote ZEB1 expression and activity in neighbouring cells, thereby fostering their malignant transformation. Collectively, our findings unveil a previously unprecedented role for senescence in bridging RAS activation and EMT over the course of malignant transformation of human mammary epithelial cells.

## Introduction

The development of cancer is viewed as an evolutionary process, in which the malignant transformation of a normal cell involves a number of limiting genetic and epigenetic events ^1^. Among the most frequently activated oncogenes in cancer are the proto-oncogenes of the RAS family (KRAS, NRAS and HRAS), found mutated in 19% of all cancers ^2^. Nevertheless, a strong heterogeneity in the incidence of RAS mutations is observed according to the type of cancer. Being most prevalent in pancreatic cancer, colorectal cancer and lung adenocarcinoma, RAS appears to be seldom mutated in liver or breast cancers. This difference in RAS mutation frequency among cancer types suggests a cellular state of predisposition to withstand RAS transformation ^3^. Interestingly, although the RAS pathway is infrequently altered in breast cancers, our recent work has revealed that activation of RAS is in fact the most common genetic alteration found specifically in the “claudin-low” (CL) molecular subtype of breast tumours ^4^. CL tumours correspond to a rare form of triple-negative tumours that are highly enriched in mesenchymal traits and stem cell features ^5–7^. We and others have recently reported the diversity of CL tumours in terms of their genomic composition, gene expression and methylation patterns as well as the repertoire of active biological pathways ^4, 8^. We have further demonstrated that this diversity reflects the impact of the cell origin on the evolutionary process of these tumours. Indeed, a subgroup of CL tumours, called CL1, designated by a low level of genomic aberrations and a prominent stemness signature, appears to develop from the malignant transformation of a normal human mammary stem cell (MaSC) that has inherent mesenchymal features, while two others subgroups, CL2 and CL3, defined by lower stemness capacity and moderate to high levels of genomic aberrations, derive from luminal or basal-like breast cancers, respectively through an epithelial-mesenchymal transition (EMT) in response to microenvironmental signals and/or the gradual acquisition of oncogenic events (Pommier et al., 2020). Notably, all 3 subgroups of CL tumours show a high frequency of activation of the RAS-dependent signalling pathway. In the context of CL1, this observation suggests that MaSCs have a high susceptibility to malignant transformation by RAS, in line with the emerging notion of cellular pliancy (Puisieux et al., 2018). In the context of CL2 and CL3, it suggests that the activation of the RAS pathway may promote cellular plasticity during tumour development, through the induction of EMT. To further study the interplay between the oncogenic activation of RAS and cellular plasticity during breast tumorigenesis, we used inducible models of human mammary epithelial cells expressing a mutated form of RAS to analyse early steps of malignant transformation. We demonstrate that RAS-dependent transformation relies on the induction of ZEB1-dependent EMT. Notably, this transdifferentiation process is driven by a paracrine signalling mechanism involving the secretion of cytokines IL-6 and IL-1α from neighbouring RAS-activated senescent cells. Hence, we unveil an unforeseen pro-tumorigenic interplay between RAS activation, oncogene-induced senescence and EMT.

## Results

### Claudin-low tumours display higher RAS pathway activation correlated with EMT signature

To confirm a link between RAS/MAPK activation and an EMT signature in breast cancers, single sample gene-set enrichment analysis (ssGSEA) scores were generated for the two pathways (RAS/MAPK and EMT) using MSigDB C2 curated gene sets (KEGG, GO, REACTOME, etc.) on breast cancer cohorts from The Cancer Genome Atlas Network (TCGA; https://www.cancer.gov/tcga) and Molecular Taxonomy of Breast Cancer International Consortium (METABRIC) ^9^ as well as on breast cancer cell lines from the Cancer Cell Line Encyclopaedia (CCLE) ^10^.

As expected, signatures of both RAS and MAPK signalling pathways were overrepresented in the CL breast cancer subtype compared to other breast cancer subtypes (Fig. 1a and Extended Data Figs. 1a and 1d). Likewise, a pan-cancer EMT signature was also clearly enriched in CL tumours (Fig.1b and Extended Data Figs. 1b, e). Notably, the RAS/MAPK pathway was highly correlated with EMT, even in the CL cell lines, excluding any stroma contamination bias (Fig. 1c and Extended Data Figs. 1c, f).

**Fig. 1.**
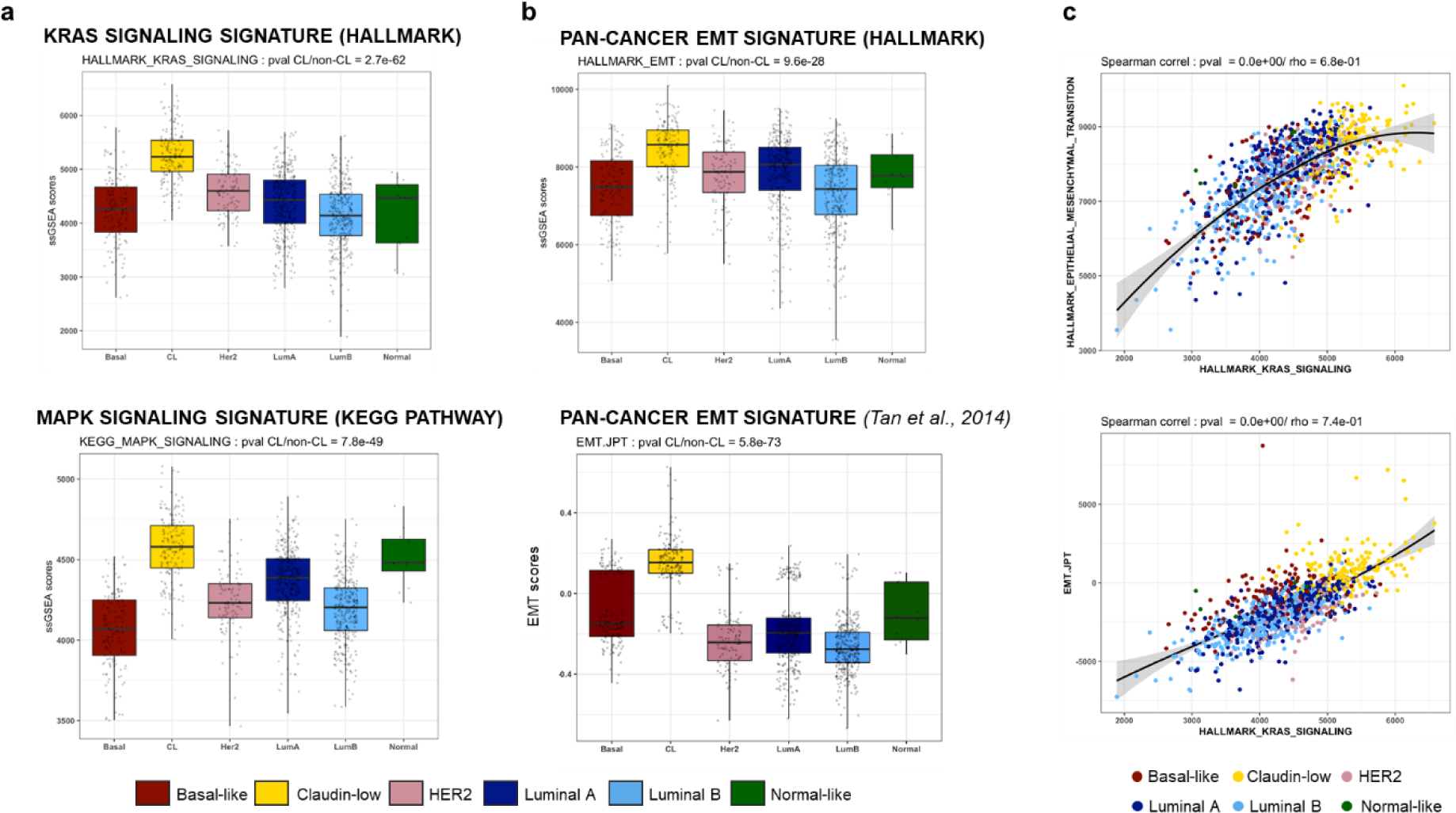
RAS/MAPK and EMT pathways are highly correlated and strongly activated in claudin-low breast cancer subtype. **a**,**b**, Distribution of ssGSEA or EMT scores for each molecular subtype of breast cancer in TCGA cohort, corresponding to (**a**) RAS or MAPK pathway (ssGSEA score) and (**b**) EMT pathway (ssGSEA or EMT score ^66^); **c**, Spearman correlations between KRAS signalling pathway (ssGSEA score) and EMT pathway (ssGSEA score or EMT score ^66^) in breast tumours from TCGA dataset. See Extended Fig.1.

### RAS-activation in differentiated mammary cells induces a phenotypic switch associated with EMT-TF expression

We wondered whether RAS activation in normal differentiated epithelial cells could induce the reactivation of EMT-inducing transcription factors (EMT-TFs), leading to the acquisition of cellular plasticity. To test this, we carried out kinetics analysis in a RAS inducible cellular model, HME-RAS_ER_, generated by introducing a 4-hydroxytamoxifen (4-OHT)-inducible form of HRAS^G12V^ into human mammary epithelial (HME) cells. Seven days into RAS activation in HME-RAS_ER_, an increase in expression of EMT-TFs ZEB1 and ZEB2 as well as a concomitant decrease in the expression of ZEB negative regulator, the miR-200 mRNA family were revealed (Fig. 2a,b and Extended Data Fig.2a). Moreover, a clear change in cellular morphology was noted at day 49 following RAS induction, with HME-RAS_ER_ cells losing their cobblestone-like epithelial morphology and gaining a spindle-like morphology suggestive of mesenchymal features associated with an EMT process (Fig. 2c). It was previously demonstrated that cell commitment to EMT was associated with the acquisition of features of cancer stem cells (CSC) ^11, 12^ ^13, 14^. Therefore, the expression of CD24, CD44, EpCAM and CD104 was evaluated by flow cytometry analysis in HME-RAS_ER_ cells ensuing RAS activation. The emergence of the three main phenotypes associated with breast CSC, namely CD24^-/low^/CD44^+^ (Fig. 2c), CD24^-/low^/CD44^+^/EpCAM^+^ and CD44^+^/CD104^+^ cells (Fig. 2d and Extended Data Fig. 2b), were observed as early as day 7 following RAS activation, suggesting the onset of a reprogramming process. These results were confirmed using two other independent HME-RAS_ER_ models (Extended Data Fig. 3a-c). The emergence of these CSC phenotypes was accompanied with an increased capacity to grow in soft agar, a gold-standard assay for cellular transformation (Fig. 2e). We next examined whether RAS activation affected the structural and morphological organization in 3D culture. Here, HME-RAS_ER_ cells were grown in 3D organoid culture conditions for 10 days, without 4-OHT activation, in order to obtain a typical organoid structure prior to RAS activation. A disruption in the structural organization associated with the acquisition of EMT features was observed in 4-OHT-induced HME-RAS_ER_ organoids compared to non-induced organoids (Fig. 2f).

**Fig. 2.**
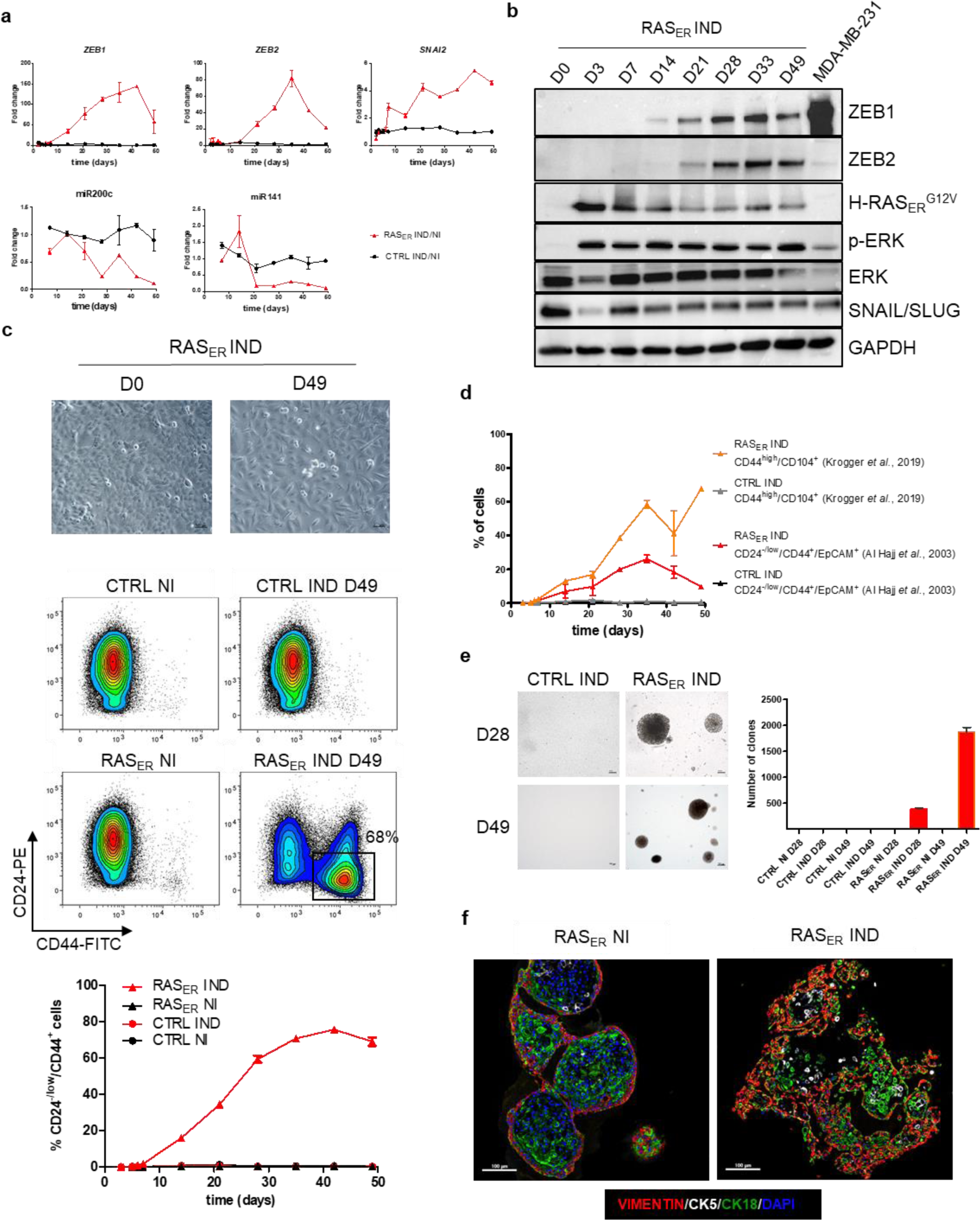
RAS-activation in differentiated mammary cells induces a phenotypic switch in 2D associated with the expression of EMT-TFs and a disruption of organoid formation in 3D. **a**, Fold change expression across time of mRNA for EMT-TFs *ZEB1*, *ZEB2* and *SNAI2*, miR200c and miR141 in HME-RAS_ER_ cells or in HME-CTRL cells induced with 4-OHT (IND) or not (NI). Relative expression was determined by the ΔΔCt method, normalized to the expression of *HPRT1* housekeeping gene for *ZEB1*, *ZEB2* and *SNAI2* mRNA and to RNU-48 for miR200c and miR141 and divided by the expression of the untreated sample at the same time point (Ratio IND/NI). Median± range (*n*=2 independent experiments in duplicate); **b**, Immunoblot showing the expression of HRAS_ER_^G12V^, pERK1/2, ERK, ZEB1, ZEB2 and SNAIL/SLUG in HME-RAS_ER_ cells (RAS_ER_) at day 0 (D0), day 3 (D3), day 7 (D7), day 14 (D14), day 21 (D21), day 28 (D28), day 33 (D33) and day 49 (D49) following 4-OHT treatment. MDA-MB 231 cell line was used as a positive control for ZEB1, ZEB2 and SNAIL/SLUG expression. GAPDH level was used as a loading control; **c**, Upper panel: Representative bright field images of HME-RAS_ER_ cells at D0 and D49 after RAS activation (Scale bars, 200 µm). Lower panel: Representative FACS analysis of CD24 and CD44 markers in HME-RAS_ER_ (RAS_ER_) cells and HME-CTRL cells (CTRL) at D0 and D49 of induction with 4-OHT (IND) or not (NI). Kinetics analysis across time of the percentage population of CD24^-/low^/CD44^+^ in induced HME-RAS_ER_ cells and induced HME-CTRL cells after 4-OHT treatment. Median± range (*n*=2 independent experiments in duplicate); **d**, Kinetics FACS analysis of cancer stem cell markers CD24^-/low^/CD44^+^/EpCAM^+^ (as described by ^13^ and CD44^+^/CD104^+^ (as described by ^14^ in induced HME-RAS_ER_ cells and HME-CTRL cells after 4-OHT treatment. Median± range (*n*=2); **e**, Transformation potential analysis by soft agar colony formation assay. Representative images of phase-contrast colonies (Scale bars, 200 µm). The number of colonies (as defined by > 20 cells) is indicated. Median± SD (*n*=3); **f**, Representative images of multi-immunofluorescence staining for the indicated markers in cross-sections of organoids generated from HME-RAS_ER_ cells (RAS_ER_) that were induced with 4-OHT (IND) or not (NI) for 21 days (Scale bars, 100 µm). (*n*=3). See Extended Data Figs. 2 and 3.

Collectively, these data suggest that RAS activation in HME cells induces EMT-TF expression and enhances cellular plasticity and acquisition of stemness markers.

### RAS-induced cellular plasticity and transforming ability are ZEB1-dependent

To determine if RAS activation could drive cellular plasticity in the absence of EMT-TFs, a vector allowing the constitutive expression of miR-200c, a major negative regulator of *ZEB1*, *ZEB2* and *SNAI1*, was introduced into HME-RAS_ER_ cells ^15, 16^ (Extended Data Fig. 4a-c). As expected, induction of ZEB1 and ZEB2 expression was dampened following RAS activation in this model (Extended Data Fig. 4d,e). miR-200c expression further prevented the emergence of CD24^-/low^/CD44^+^ cells and the acquisition of properties associated with transformation and stemness (Fig. 3a-c). Indeed, induced HME-RAS_ER_–miR-200c cells were neither able to form colonies in soft-agar assays nor mammospheres in low-adherent culture conditions (Fig. 3b,c). Hence, inhibiting the expression of ZEB1 and ZEB2 by overexpressing miR-200c prevented the acquisition of cellular plasticity following RAS activation.

**Fig. 3.**
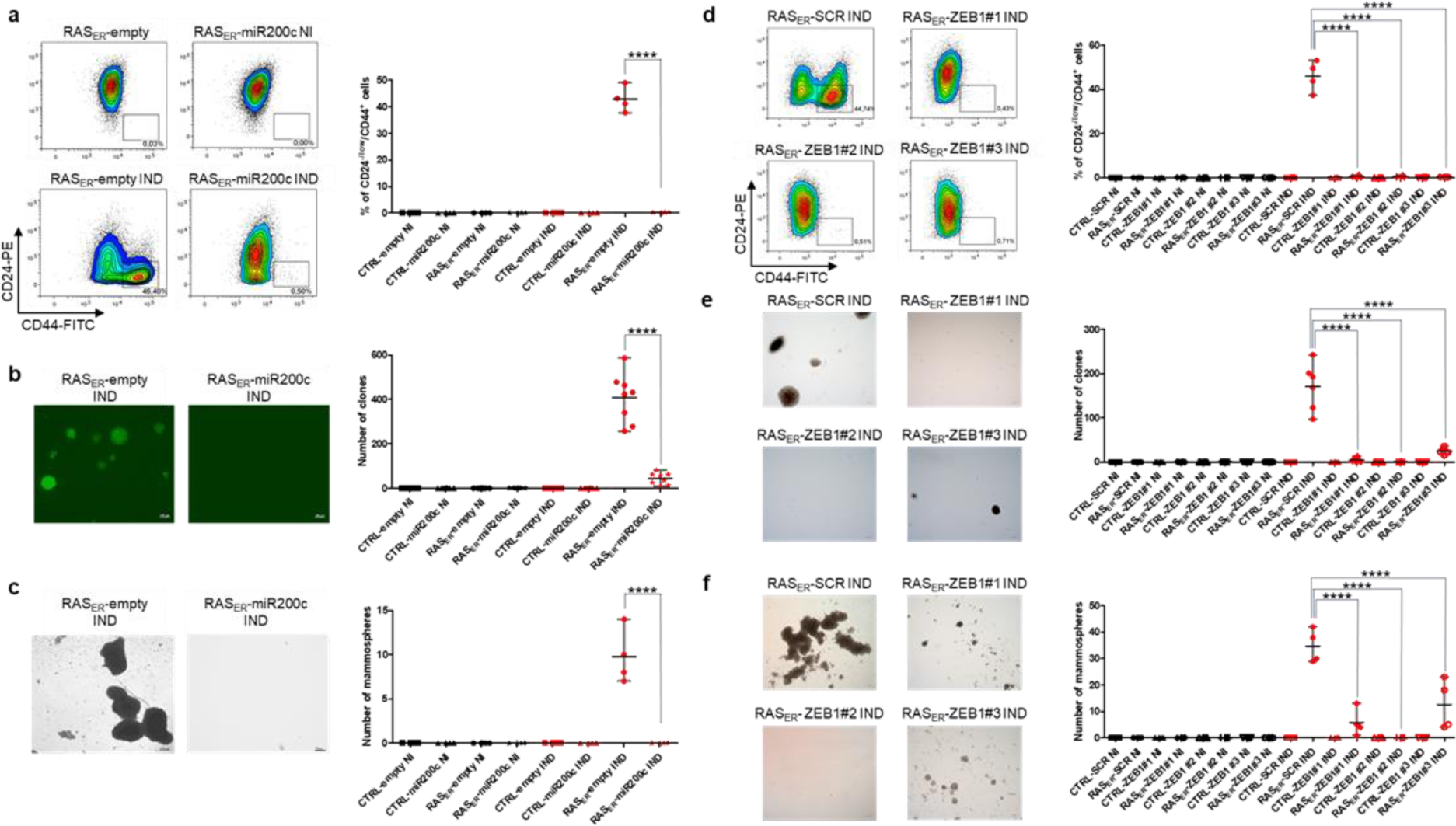
RAS-induced cellular plasticity and transformation capabilities are ZEB1-dependent. **a-c**, HME-RAS_ER_-miR200c cells (RAS_ER_-miR200c) or HME-RAS_ER_-empty cells (RAS_ER_-empty) after 28 days of induction with 4-OHT (IND) or not (NI); **d-f**, HME-RAS_ER_- CRISPR ZEB1 clones (RAS_ER_-ZEB1#1, RAS_ER_-ZEB1#2 and RAS_ER_-ZEB1#3) or HME-RAS_ER_-CRISPR scramble (RAS_ER_-SCR) cells after 28 days of 4-OHT treatment (IND) or not (NI); **a**,**d** Flow cytometry analysis and quantification of CD24^-/low^/CD44^+^ cells. Data are presented as median± range of four independent experiments (*n=4*); **b,e** Transformation potential analysis by soft agar colony formation assay. The number of colonies (as defined by > 20 cells) is indicated. Data are presented as median± range of eight independent experiments (n=8) (**b**) or six independent experiments (n=6) (**e**); Images of GFP-positive colonies (**b**) or phase-contrast colonies (**e**). (Scale bars, 200 µm); **c**,**f**, Quantification and phase-contrast images of mammospheres. (Scale bars, 100 µm (**c**) and 200µm (**f**)). Data are presented as median ± range of four independent experiments (*n=4*). p values are calculated by one-way ANOVA, Tukey multiple comparison test (****p<0.001). See Extended Data Fig. 4.

Because ZEB1 is a known master regulator of EMT in mammary epithelial cells, a CRISPR/Cas9 knock-down approach was used to determine its role in the context of RAS activation. Three independent *ZEB1*-inactivated clones of HME-RAS_ER_ cells were generated (HME-RAS_ER_-CRISPR ZEB1#1, HME-RAS_ER_-CRISPR ZEB1#2 and HME-RAS_ER_-CRISPR ZEB1#3) (Extended Data Fig. 4f,g). *ZEB1* inactivation alone inhibited the emergence of the CD24^-/low^/CD44^+^ population as well as the acquisition of transformation and stemness properties (Fig. 3d-f). Interestingly, *ZEB2* and *SNAI2* expression was moderate in *ZEB1*-inactivated clones, suggesting that their up-regulation was essentially ZEB1-dependent (Figure S4H).

Altogether, these data demonstrate that plasticity features acquired by differentiated HME cells following RAS activation strongly rely on ZEB1.

### Dual fate following RAS activation: senescence or mesenchymal transition

Because only a sub-population of induced HME-RAS_ER_ cells displays EMT features after RAS activation, we performed single-cell RNA sequencing (scRNAseq) analysis at different time points (D0, D3, D7, D14 and D20) with the aim of determining the mechanisms underlying cellular plasticity. Expression profiles were obtained for 12,357 genes in 445 cells (117 cells at D0, 83 cells at D3, 74 cells at D7, 82 cells at D14 and 89 cells at D20). To analyse transcriptional variability over the complete scRNAseq dataset, dimensions were reduced using Principal Component Analysis (PCA) and Uniform Manifold Approximation and Projection (UMAP) ^17^ (Fig. 4a). A dimension reduction analysis and graph-based k-means clustering identified 5 clusters based on their differential gene expression, numbered 1 to 5 (Fig. Extended Data Fig. 5a). For each cluster, specific marker genes were extracted to search for enriched pathways using MSigDB (Supplementary Tables 1-5). This allowed us to functionally annotate each cluster (Fig. 4b), resulting in clusters 1, 2 and 4 related to cell cycle phases (cluster1 to G1 phase, cluster 2 to G2/M phase and cluster 4 to S phase) and clusters 3 and 5 related to senescence and EMT, respectively (Fig. 4b,c). Interestingly, cluster 3, which represents less than 10% of cells at D0, constitutes more than 40% of cells at D7 while cluster 5 appears at D3 and is the most represented at D20 (Fig. 4d). We next interrogated a possible link between these two emerging clusters through evolutionary trajectory analysis. Given the uncertain origin of a potential trajectory, cluster 1 (related to G1 phase) was used as a starting point since it was the most represented at D0 (40%) (Fig. 4d). Trajectory analysis showed two different end points: cluster 3/senescence or cluster 5/EMT (Fig. 4e), suggesting that, following RAS activation, cells would either undergo senescence or acquire a mesenchymal phenotype. Trajectory analysis using cluster 2 or 4 as a starting point gave identical results with regards to the evolutionary outcome: senescence or EMT (Extended Data Fig. 5b). Interestingly, the rise of the senescence cluster preceded the rise of the EMT cluster (D7 vs D20, respectively; Fig. 4d), indicating a possible underlying causal relationship.

**Fig. 4.**
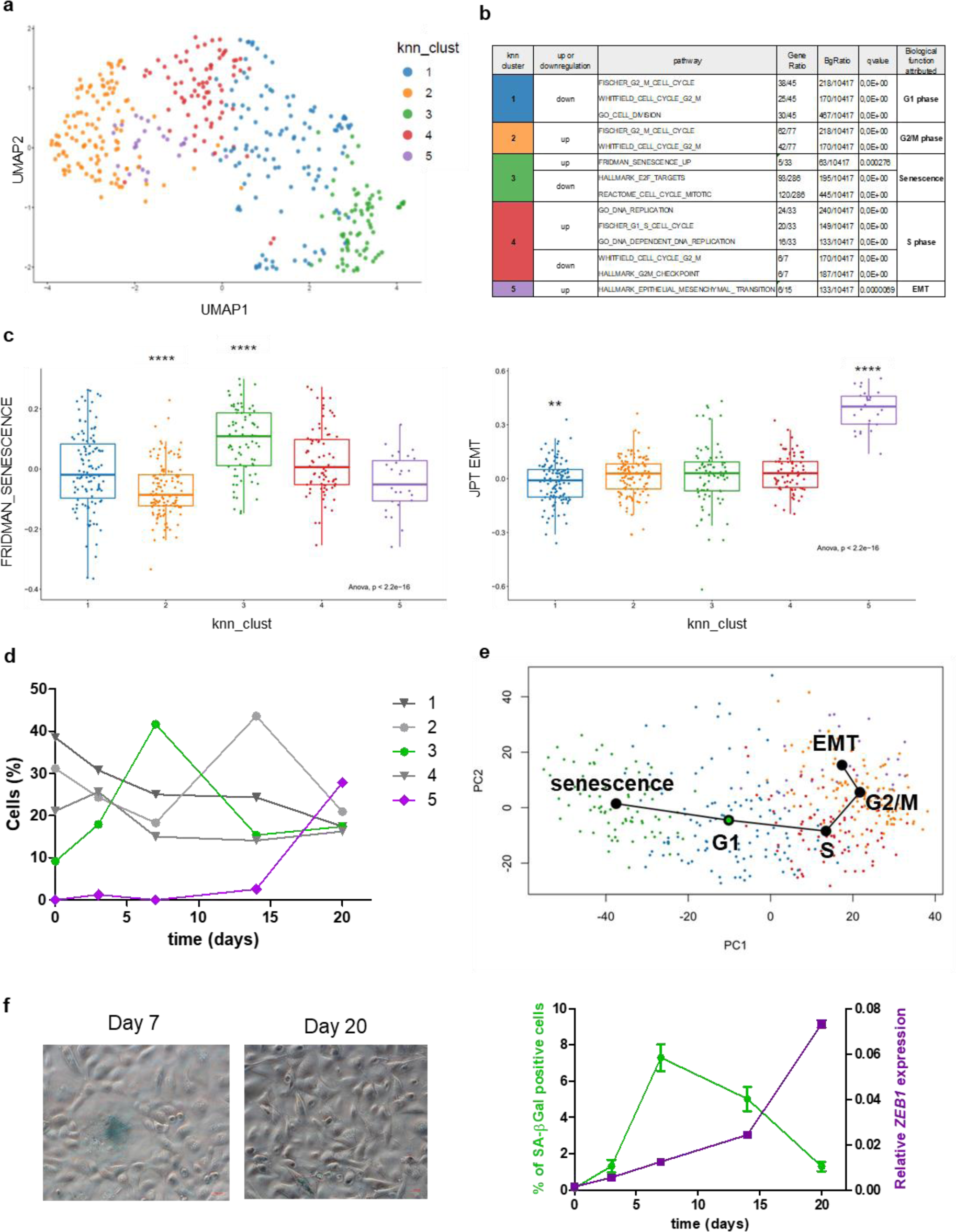
Identification of distinct senescence and EMT clusters in induced HME-RAS_ER_. **a**, Unsupervised UMAP of the transcriptome of all cells at all time points (D0, D3, D7, D14 and D20). Cells are colored by their attributed cluster; **b**, Main altered pathways by marker genes for each cluster. Gene ratio presented as k/n where k is the size of the overlap of our input with the specific gene set and n is the size of the overlap of our input with all members of the collection of gene sets. qvalue refers to false discovery rate; **c**, Scores per cell for two transcriptomic pathways: FRIDMAN_SENESCENCE (ssgsea score) and JPT_EMT (EMT cell line score from ^66^. Cells are grouped by clusters, each box representing the median and interquartile ranges. Individual Wilcoxon tests, p-value is represented by stars (** p<=0.01 and ****p<=0.0001). All p-values were corrected by the Holm-Bonferroni method; **d**, Proportion of cells by cluster at each time point. Emerging clusters across time were colored (cluster 3 in green and cluster 5 in purple); **e**, Projection of the trajectory analysis performed by the Slingshot algorithm along the first two axes of an unsupervised PCA of all cells. Cells are colored by clusters. Starting point of the trajectory analysis is cluster 1 (green circle); **f**, Left panel: Representative images of senescence-associated β-galactosidase (SA-β-gal) activity in induced HME-RAS_ER_ cells at day 7 and day 20. (Scale bars, 200 µm); Righ panel: Percentage of SA-β-gal-positive cells (green) and of *ZEB1* mRNA expression (purple) across time after RAS activation in induced HME-RAS_ER_ cells. Median± range (*n*=3). See Extended Data Fig. 5.

### ZEB1-dependent plasticity is driven by oncogene-induced senescence

Based on scRNAseq data, we sought to gain further insights into a potential crosstalk between oncogene-induced senescence (OIS) and EMT. We therefore analysed the emergence of senescent cells, determined by SA-β-gal positive staining and ZEB1 expression in a time dependent manner. We previously showed that ZEB1 expression distinctly increased from D14 onwards and was directly associated with the emergence of the CD24^-/low^/CD44^+^ population (Fig. 2a-d). As shown in Fig. 4f, RAS-induced senescent cells appeared from D3 with a peak at D7.

The initial emergence of OIS followed by the delayed occurrence of EMT led us to hypothesize that senescence may contribute to the induction of cellular plasticity. As a first approach to test this hypothesis, HME-RAS_ER_ cells were treated with dasatinib, an inhibitor of the Bcr-Abl protein kinase and a known senolytic drug ^18, 19^. Dasatinib treatment significantly reduced the number of senescent cells at day 7 of RAS-activation, along with a 10-fold decrease in SA-β-gal-positive cells (Fig. 5a). Notably, this senescence blockage was also associated with a decrease in both *ZEB1* expression and the percentage of CD24^-/low^/CD44^+^ cells at day 21 (Fig. 5b,c) in comparison to DMSO-treated cells, thus in support of ZEB1-dependent cellular plasticity being reliant on the onset of senescence.

**Fig. 5.**
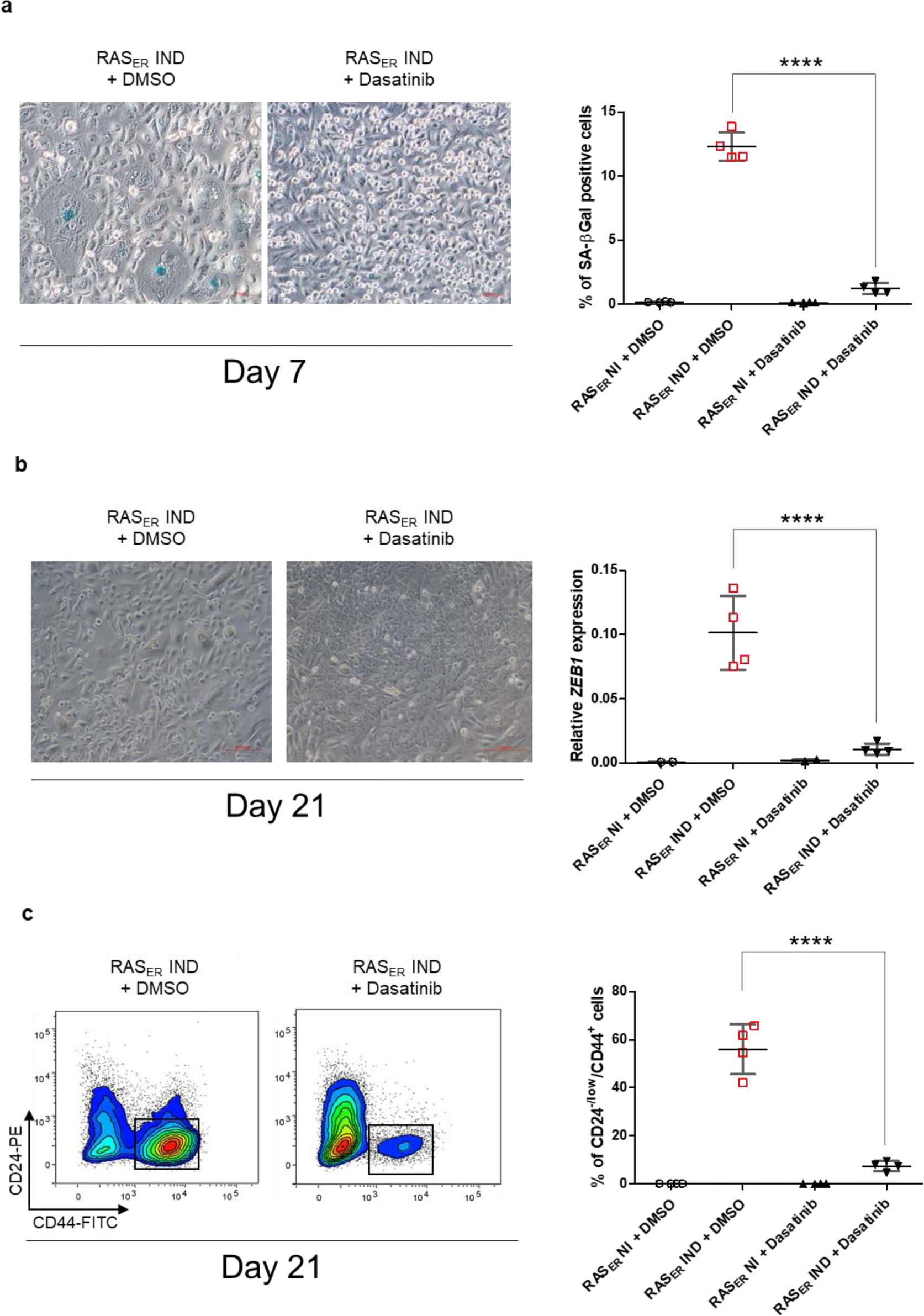
Senescent cells drive ZEB1-dependent cellular plasticity. **a**, Representative phase-contrast images and quantification of senescence-associated β-galactosidase (SA-β-gal) activity in HME-RAS_ER_ (RAS_ER_) cells analyzed at D7 of induction with 4-OHT (IND) or not (NI) and treatment with the senolytic drug dasatinib at 1.5µM or the equivalent concentration of DMSO starting from D0. Median±range (*n=4*); **b**, Phase-contrast images and *ZEB1* mRNA expression in HME-RAS_ER_ (RAS_ER_) cells at D21 of induction with 4-OHT (IND) or not (NI) and treatment with the senolytic drug dasatinib at 1.5µM or the equivalent concentration of DMSO starting from D0. Median±range (*n=4*); **c**, Representative flow cytometry analysis on CD24 and CD44 markers. Quantification of CD24^-/low^/CD44^+^ cells in HME-RAS_ER_ (RAS_ER_) cells at D21 of induction with 4-OHT (IND) or not (NI) and treatment with the senolytic drug dasatinib at 1.5µM or the equivalent concentration of DMSO starting from D0. Median±range (*n=4*). p values are calculated by one-way ANOVA, Tukey multiple comparison test (****p<0.001).

### ZEB1-dependent cellular plasticity is driven by factors secreted by senescent cells

Upon entering senescence, cells produce a set of pro-inflammatory cytokines and chemokines, extracellular matrix proteins, growth factors and metalloproteinases composing the senescence-associated secretory phenotype (SASP), which through secretory signalling, impacts the dedifferentiation of somatic cells ^20^. We therefore questioned if a secretory signalling process emanating from RAS-activated senescent cells may be responsible for ZEB1-mediated plasticity in neighbouring differentiated cells. To explore this hypothesis, we generated a cellular HME model (HME_d2GFP200) that expressed a destabilized GFP (d2GFP) including a 3′UTR containing five miR-200 target sequences (Extended Data Fig. 6a). This set-up was expected to highlight cells undergoing EMT through an increased GFP signal underlying lower levels of expression of miR-200 family members ^21^. To determine whether senescent cells promote EMT in neighbouring cells, RAS-induced HME-RAS_ER_ cells tagged with dsRED (HME-RAS_ER_-dsRED cells) were co-cultured with HME_d2GFP200 cells (Fig. 6a). An enrichment of GFP^high^ cells associated with an increase in ZEB1 expression and an emergence of a CD24^-/low^/CD44^+^ phenotype was observed in HME_d2GFP200 cells (Fig. 6b-d and Extended Data Fig. 6b), revealing that ZEB1-reactivation can be driven by secreted factors. Moreover, these co-cultured HME_d2GFP200 cells were able to form colonies in soft agar assay, suggesting that they had acquired cellular properties associated with transformation (Fig. 6e). Finally, we performed a transwell co-culture assay using HME_d2GFP200 cells and RAS-induced HME-RAS_ER_-dsRED cells. The characterization of cells following co-culture confirmed the enrichment of a CD24^-/low^/CD44^+^ phenotype in GFP^high^ cells (Fig. 6f,g), thus evoking either an autocrine or paracrine signalling process rather than a juxtacrine one.

**Fig. 6.**
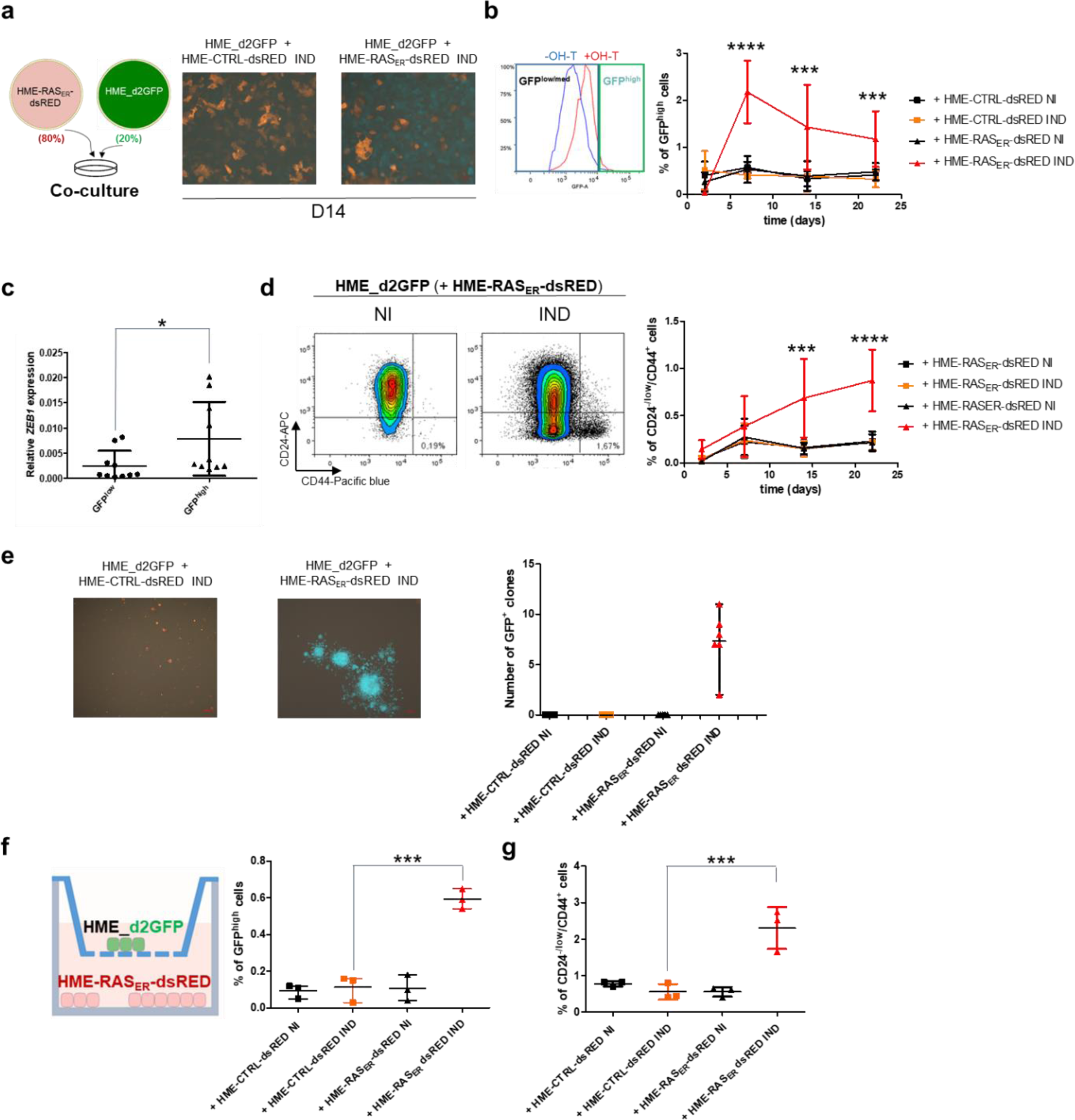
ZEB1-dependent cellular plasticity is driven by cytokines secreted by senescent cells. **a-e**, Co-culture assays; (**a**) Experimental outline and representative fluorescence images of co-culture assay; (**b**) Representative flow cytometry analysis of GFP expression in HME_d2GFP cells co-cultured with HME-RAS_ER_-dsRED (+HME-RAS_ER_-dsRED) cells after 14 days of induction with 4-OHT (IND) or not (NI) and quantification of GFP^high^ population across time. Median±range (n=7); (**c**) *ZEB1* expression in HME-d2GFP cells co-cultured with HME-RAS_ER_-dsRED cells after induction with 4-OHT and sorted at day 14 based on their GFP expression. Median±range (*n*=5 independent experiences in duplicate); (**d**) Representative flow cytometry analysis of CD24 and CD44 markers in HME_d2GFP cells co-cultured with HME-RAS_ER_-dsRED or HME-CTRL-dsRED cells, induced with 4-OHT (IND) or not (NI) for 21 days and analyzed at days 2, 7, 14, 22; (**e**) Transformation potential analysis as assessed by a soft agar colony formation assay of HME_d2GFP cells co-cultured with HME-RAS_ER_-dsRED (+HME-RAS_ER_-dsRED) or HME-CTRL-dsRED (+HME-CTRL-dsRED) cells, induced with 4-OHT (IND) or not (NI) for 21 days. Representative images and quantification of GFP-positive colonies (as defined by > 20 cells). Median±range (*n*=6); **f**, **g**, Transwell culture assays; (**f**) Quantification of GFP^high^ population from HME_d2GFP cells co-cultured with HME-RAS_ER_-dsRED (+HME-RAS_ER_-dsRED) or HME-CTRL-dsRED (+HME-CTRL-dsRED) cells, induced with 4-OHT (IND) or not (NI) for 21 days. Median±range (*n*=3); (**g**) Quantification of CD24^-/low^/CD44^+^ population from HME_d2GFP cells co-cultured with HME-RAS_ER_-dsRED (+HME-RAS_ER_-dsRED) or HME-CTRL-dsRED (+HME-CTRL-dsRED) cells, induced with 4-OHT (IND) or not (NI) for 21 days. Median±range (*n*=3). p values are calculated by one-way ANOVA, Tukey multiple comparison test (****p<0.001). See Extended Data Fig. 6.

### IL-6 and IL-1α secreted by senescent cells following RAS activation promote ZEB1-induced cellular plasticity

Given that our data revealed a possible effect of secretory signalling on the acquisition of ZEB1-dependent cellular plasticity following RAS activation, we next aimed to narrow down the specific factors produced by senescent cells leading to this effect, focusing specifically on SASP constituents. Multiplex technology was used to quantify the concentrations of SASP factors in RAS-induced HME-RAS_ER_ cell culture supernatants. Quantifications revealed high levels of cytokines IL-6, IL-8, IL-1α, IL-1β and granulocyte-macrophage colony-stimulating factor (GM-CSF) (Fig. 7a). Experiments based on the direct addition of these cytokines to HME cells at the highest concentration found in cell culture supernatants showed the ability of IL-6, IL-1α and IL-1β to promote the acquisition of a plasticity phenotype (Fig. 7b). Interestingly, the combination of IL-6 and IL-1α had a cumulative effect on the emergence of CD24^-/low^/CD44^+^ cells. Next, to evaluate the role of these cytokines in RAS-induced plasticity, we used a depletive approach with cytokine-neutralizing antibodies or IL-6R antibody (tocilizumab). Double inhibition of IL-6/IL-6R or the inhibition of IL-1α alone decreased the CD24^-/low^/CD44^+^ population and *ZEB1* expression as well as the transforming abilities of RAS-induced HME-RAS_ER_ cells (Fig. 7c,d and Extended Data Fig.7a). Strikingly, combination treatments with different cytokines, showed that the concomitant inhibition of IL-6/IL-6R and IL-1α induced an almost 10-fold decrease in cellular plasticity and transformation capacities, suggesting that these two cytokines are major determinants of ZEB1-mediated plasticity and transformation (Fig. 7c,d).

**Fig. 7.**
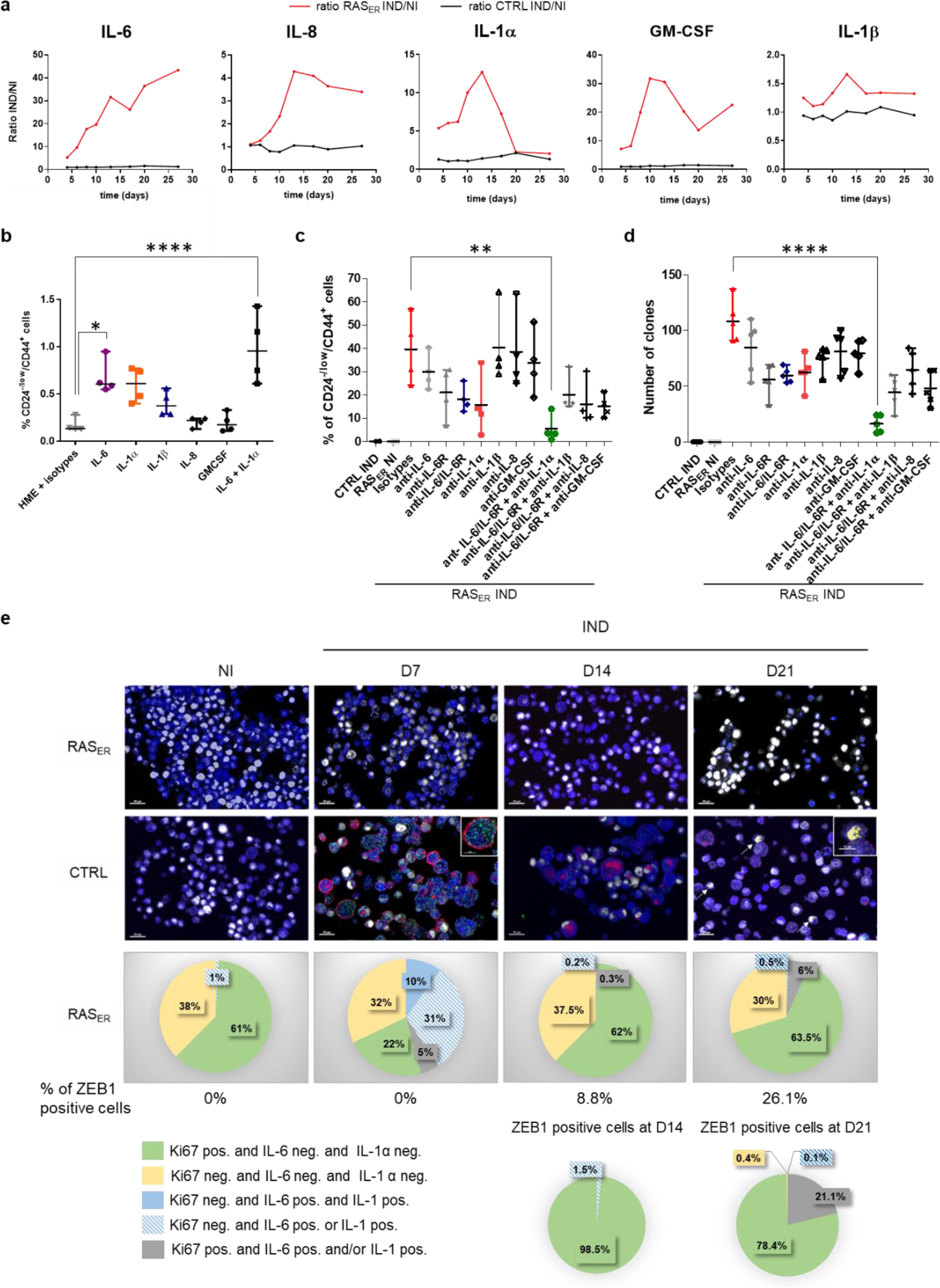
IL-6 and IL-1α are major cytokines responsible for ZEB1 cellular plasticity. **a**, Quantification of cytokine levels in the supernatant from HME-RAS_ER_ cells after 4-OHT treatment compared to controls at days 4, 6, 8 10, 13, 17, 20 and 27. Data are presented as fold change of secreted cytokine concentrations in HME-RAS_ER_ or HME-CTRL cells induced by 4-OHT (IND) or not (NI); (**b**) Quantification by flow cytometry analysis of CD24^-/low^/CD44^+^ population of HME cells after treatment by the indicated cytokine(s) or isotype controls analyzed at day 40. Median±range (*n*=4); (**c**) Quantification by flow cytometry analysis of CD24^-/low^/CD44^+^ population of HME-CTRL or HME-RAS_ER_ cells (RAS_ER_) induced by 4-OHT (IND) or not (NI) and treated by neutralizing antibodies or isotype controls analyzed at day 28. Median±range (*n*=4); (**d**) Transformation potential analysis by soft agar colony formation assay in HME-CTRL and HME-RAS_ER_ cells (RAS_ER_) induced by 4-OHT (IND) or not (NI) and treated by neutralizing antibodies or isotype controls analyzed at day 28. Median±range (*n*=4); (**e**) Upper panel: Multi-fluorescence staining kinetics. HME-RAS_ER_ (RAS_ER_) and HME-CTRL (CTRL) cells were stained with anti-Ki67 (white), anti-IL-6 (green), anti-IL-1α (red) and anti-ZEB1 (yellow) following induction by 4-OHT (IND) at days 7, 14 and 21 or not (NI). Representative pictures are shown. Scale 50µm. Lower panel: Graphical representation of cellular phenotype distribution according to Ki67, IL-6, IL-1α or ZEB1 staining. See Extended Data Fig.7.

### ZEB1-induced cellular plasticity relies on a paracrine effect mediated by senescent cells

To further characterize the process involved in ZEB1-mediated plasticity after RAS activation, we used multi-fluorescent labelling kinetics to visualize the proliferative state of cells via anti-Ki67, the expression of cytokines IL-6 and IL-1α, and ZEB1 in HME-RAS_ER_ cells (Fig. 7e). We observed that at day 7 of RAS activation, 10% of RAS-induced HME-RAS_ER_ cells showed a negative Ki67 profile and were double positive for IL-6 and IL-1α, consistent with a senescent state. Moreover, 31% of cells presented a negative Ki67 profile and were positive for either IL-6 or IL-1α, potentially indicating a pre-senescent state. None of these cells were positive for ZEB1. Fourteen days after RAS induction, 9% of cells were ZEB1 positive and less than 2% of them were both negative for Ki67 and positive for IL-6 and/or IL-1α (Fig. 7e). At day 21, 26% of cells were ZEB1 positive with the majority being Ki67 positive. Overall, these data suggest that cells entering OIS do not express ZEB1, consistent with the hypothesis of the involvement of a paracrine process driving EMT in neighbouring cells. Next, we co-cultured HME-d2GFP_200 cells with tamoxifen-induced HME-RAS_ER_-dsRED cells and treated them with IL-6/IL-6R and IL-1α neutralizing antibodies. The acquisition of cellular plasticity in HME-d2GFP_200 cells was strongly impaired following treatment (Extended Data Fig. 7b, c). Altogether, these results reveal that ZEB1 activation after RAS induction is driven by a paracrine process involving the secretion of IL-6 and IL-1α cytokines from senescent cells.

## Discussion

Despite their relative rarity, breast cancers of the claudin-low molecular subtype are a remarkable illustration of the influence of the cell of origin and the microenvironment on tumour development and progression. Indeed, their molecular characterization has recently revealed three main subgroups emerging from unique evolutionary processes ^4, 8^. The subgroup 1 of claudin-low tumours (CL1), characterized by stem cell features and low genomic instability, are presumed to develop from the malignant transformation of a normal mammary stem cell, whereas claudin-low subtypes 2 (CL2) and 3 (CL3) appear to evolve from luminal or basal tumours, respectively following aberrant induction of an EMT process in response to microenvironmental cues. Interestingly, all three subgroups are characterized by a high frequency of activation of the RAS-MAPK pathway, suggesting the existence of a mechanistic link between the activation of this oncogenic pathway and the mesenchymal characteristics of these tumours. To characterize this interplay, we have developed an *in vitro* approach for identifying the early and long-term consequences of RAS pathway activation in human mammary epithelial cells. Our observations show a causal link between the activation of the RAS-MAPK pathway and the induction of EMT, the nature of which is totally distinct from the usual view of the involvement of EMT in cancer, in terms of the phases of its involvement, the mechanisms underlying its induction, and its consequences during tumorigenesis.

In cancer, EMT is generally considered to be a process induced during tumour progression in response to microenvironmental signals such as growth factor signalling, hypoxia and immune response pathways ^22^, which will promote key steps of the invasion-metastasis cascade ^23^. Here, we unveil a causal link between RAS activation and EMT. In this context, inhibition of EMT prevents RAS-induced transformation of human mammary epithelial cells, suggesting a determinant role for this transdifferentiation process in early phases of tumorigenesis following an oncogenic event. It remains to be determined whether this notion is generalizable to other tumour types where activation of the RAS pathway is a frequent and an early event. However, it is noteworthy that in a mouse model of pancreatic cancer driven by Pdx1-cre-mediated activation of mutant Kras, homozygous deletion of *Zeb1* significantly reduces features of acinar-ductal metaplasia (ADM) as well as the number and grading of PanIN, the main origins of pancreatic precursor lesions ^24, 25^. The causal link between RAS pathway activation and EMT induction further provides a mechanistic explanation for the characteristics of CL2 and CL3 breast tumours since RAS-MAPK activation during the evolution of their originating luminal or basal tumours would indeed lead to the acquisition of mesenchymal characteristics inherent to CL subtypes. In support of this hypothesis, in transgenic mouse models, Kras activation in luminal mammary epithelial cells leads to EMT commitment and promotes claudin-low tumours ^26^. Collectively, these observations suggest that the induction of EMT during tumour development is not only a late consequence of exposure to microenvironmental signals that favor the metastatic dissemination of cancer cells, but may also constitute an early event of cellular adaptation following certain oncogenic insults, influencing tumorigenesis as well as tumour evolution and progression.

A critical finding of our study is the identification of a mechanistic link between oncogene-induced senescence and EMT over the course of malignant transformation. Consistent with our data, recurrent signatures associated with EMT and senescence have been recently reported by using a pan-cancer single-cell RNA-seq approach ^27, 28^. Cellular senescence is a stress response that is generally considered to irreversibly and stably arrest the cell cycle following a variety of intrinsic and extrinsic stressors such as shortening of chromosomal termini, DNA damage, oxidative stress, oncogenic insults or inactivation of tumour suppressors ^29^. Notably, oncogene-induced senescence or senescence induced by loss of a tumour suppressor was verified *in vivo* in human and murine preneoplastic lesions ^30–32^. Senescence is traditionally described as an innate anti-cancer mechanism since cells harbouring oncogenic mutations are prevented from proliferating and transforming ^33^. Moreover, in addition to decreasing replicative capacity, activation of senescence in different contexts and tissues leads to increased expression of inflammatory cytokines that elicit immune-mediated tumour clearance ^34, 35^. However, recent studies challenge this conventional view, showing a paradoxical role for senescence emanating from its special secretory profile that can counterintuitively promote cancer stemness and aggressiveness ^36, 37^. In our study, single cell analysis revealed oncogene-induced senescence in a subset of RAS-activated cells and the subsequent induction of an EMT process in a separate subset. Senolytic treatment showed that senescent cells comprising the first subset play a causal role in the acquisition of plasticity by neighbouring cells. This duality in cellular outcomes recalls the work of the groups of Manuel Serrano and Han Li who reported that the activation of the 4 transcription factors Oct4, Sox2, Klf4, and c-Myc (OSKM) in mouse tissue triggers opposite cellular fates, namely, senescence and reprogramming, that coexist *in vivo* within separate subpopulations of cells ^38, 39^. Consistent with a senescence response, overexpression of OSKM in human primary fibroblasts triggers replication stress and DNA damage characterized by the up-regulation of p53, p16^INK4a,^ and p21^CIP1^, impaired proliferation and formation of senescence-associated heterochromatin foci (SAHF) ^40^. With striking conceptual similarity, we have thus demonstrated that entry into senescence of a subset of mammary epithelial cells facing RAS-dependent oncogenic insult promotes the transdifferentiation of neighbouring cells.

Importantly, cellular senescence creates a tissue context that favours OSKM-driven reprogramming in surrounding cells through the paracrine action of the SASP, with IL-6 being a key mediator ^38, 39^. A similar interplay also occurs in tissue aging and injury, where there is an accumulation of senescent cells that secrete pro-inflammatory cytokines, triggering OSKM-driven dedifferentiation and reprogramming in neighbouring cells. Quantification of a series of SASP constituents and functional assays enabled us to determine the factors inherent to senescent mammary epithelial cells that are responsible for triggering a similar reprogramming process in the cellular vicinity. Namely, we identified proinflammatory cytokines IL-6 and IL-1α as major mediators of senescence-induced EMT. Of note, IL-6 has previously been shown to trigger EMT *via* JAK-STAT3 or NF-κB signalling, leading to the activation of EMT-TFs ^41–48^. Accordingly, IL-6 overexpression and/or hyper-activation has been reported in several human cancers ^49–52^. Relatively fewer studies have directly examined the role of IL-1α in cellular reprogramming and in cancer development and progression. Nevertheless, IL-1α inactivation in pancreatic cancer impaired tumour progression and immune cell infiltration ^53^. The oncogenic activation of KRas in IL-1α absence also led to a constant number of benign lesions, but above all, a decrease in dysplastic and neoplastic lesions ^53^. Finally, a recent work examining the relationship between HER2 expression in breast cancer, inflammation and expansion of cells presenting stemness properties highlighted an essential role for IL-1α ^54^. In light of these observations, further investigations will now be necessary to define the steps downstream of IL-6 and IL-1α secretion that led to the induction of ZEB1 expression.

We have previously shown in the model of breast tumorigenesis that EMT-TFs of the TWIST and ZEB families cooperate with mitogenic oncoproteins for malignant transformation by alleviating oncogene-induced senescence ^6, 55^. The present data suggest that oncogene-induced senescence of only a subset of cells is enough to trigger the bypass of this same oncosuppressive barrier in neighbouring cells via the action of released SASP factors, thereby favouring their malignant transformation. From a deterministic point of view, we are confronted here with a paradox: the act of self-sacrifice cannot be more beautifully described than by the selfless attempt of cells to undergo senescence in order to protect an organism as a whole; yet, this desperate act may be the critical element that promotes the transformation of neighbouring cells facing the same oncogenic stress. This notion supports the hypothesis that oncogene-induced senescence has contrasting cell-autonomous and non-cell autonomous effects in early tumorigenesis, providing novel clues to the ongoing debate on its tumour suppressor and pro-tumorigenic effects.

## Methods

### Cell culture

Primary human mammary epithelial cells (HMECs) (Lonza) were immortalized by hTERT, and named HME thereafter ^6^. HME-derived cells (HME-RAS_ER_, HME-CTRL, HME-RAS_ER_–miR-200c, HME-RAS_ER_–empty, HME-RAS_ER_-CRISPR ZEB1, HME-RAS_ER_-CRISPR SCR and HME_d2GFP200) were cultured in Dulbecco’s Modified Eagle’s Medium (DMEM)/HAMF12 medium with 1 % glutamax (Thermo Fisher Scientific) supplemented with 10% heat-inactivated fetal calf serum (Eurobio Scientific), 100 UI/mL penicillin, 100 µg/mL streptomycin (Thermo Fisher Scientific), 10 ng/ml human epidermal growth factor (EGF) (Peprotech), 0.5 mg/ml hydrocortisone (Sigma Aldrich) and 10 mg/ml insulin (Novorapid®, Novonordisk), at 37°C in 5% CO_2_/95% air. Breast cancer cell lines were obtained from the American Tissue Culture Collection (ATCC). MDA-MB-231, Hs578T and HEK293T cells were maintained in DMEM with 1 % glutamax (Thermo Fisher Scientific) with 10% heat-inactivated fetal calf serum (Eurobio Scientific) and 100 UI/mL penicillin, 100 µg/mL streptomycin (Thermo Fisher Scientific). All cell lines were regularly tested negative using the MycoAlert mycoplasma detection kit (Lonza).

### Lentiviral and retroviral infections

Lentiviral particles were produced using non-confluent 2x10^6^ HEK293T cells transfected with GeneJuice® Transfection Reagent (MerckMillipore) with 26 µg of total lentiviral expression vectors (10.2 µg of pCMVdeltaR8.91, 2.6 µg phCMVG-VSVG and 13.2µg of plasmid of interest). The pCMVdelteR8.91 and phCMVG-VSVG were gifts from D. Nègre (International Center for Infectiology Research, INSERM U1111–CNRS UMR5308–ENS de Lyon–UCB Lyon1, EVIR Team, Lyon, France). 48 hours post-transfection, the supernatant was collected, filtered and supplemented with 5 µg/ml of polybrene (MerckMillipore) combined with cells for 6 hours. Cell selection was done 72 hours post-infection through FACS-sorting for GFP plasmids (pCDH-CMV-EF1-copGFP/MIR200c, pCDH-CMV-MCS-EF1-copGFP-empty and FUGW-d2GFP-200) or DSRED plasmid (pLLRSV Red-empty) or by antibiotics (puromycin 0.5 mg/mL or neomycin 100 mg/mL; Invivogen) depending on plasmid construct. To produce retrovirus particles, 2x10^6^ Phoenix cells were transfected with GeneJuice® Transfection Reagent (MerckMillipore) and 15 µg of retroviral expression vectors ^56^. At 48 hours post-transfection, the supernatant was collected, filtered, supplemented with 5 µg/ml of polybrene (MerckMillipore) and combined with 10^6^ targeted cells for 6 hours. Cells were selected 48 hours following infection with puromycin (0.5 mg/ml) and/or neomycin (100 mg/ml).

### Plasmids

The lentiviral plasmids: pCDH-CMV-EF1-copGFP/MIR200c and pCDH-CMV-MCS-EF1-copGFP-empty were kind gifts from Dr. Thomas Brabletz, DsRED plasmid, pLLRSV Red-empty was a kind gift from Dr Christophe Ginestier, and miR-200 sensor plasmid, FUGW-d2GFP-200 was purchased from addgene (plasmid #79602) ^21^. Retroviral plasmids used were purchased from addgene: pLNCX2 neo-RAS-ER (#67844) and pLNCX2 neo-empty ^57,58^.

### CRISPR-Cas9 knockdown

ZEB1-depletion model in HME (HME-RAS_ER_-CRISPR ZEB1) was generated using CRISPR-cas9 gene editing technology. Scrambled sgRNA/Cas9 All-in-One Lentivector (Applied Biological Materials; Cat# K010) and ZEB1 sgRNA/Cas9 All-in-One Lentivector (Human) (Target 1: 5’-CACCTGAAGAGGACCAG-3’) (Applied Biological Materials; Cat# K2671006) lentiviral particles were used to infect HME cells. Scrambled sgRNA/Cas9 and ZEB1 sgRNA/Cas9 cells were selected with puromycin (Invivogen) at 1 µg/ml 48 hours after infection. After cloning by limited dilution, single cells were grown for approximatively 3 weeks and colonies were screened for knockouts by quantitative PCR, genomic DNA sequencing and western blotting. Genomic DNA sequencing was performed using the sanger method with the following primers for amplification and sequencing 5’-TGAACTGAACGTCAGAGTGGT-3’ (forward) and 5’TCACGTGCAGTGGCATTACT-3’ (reverse). Three different clones were finally validated.

### Kinetics, co-culture and transwell assay

For kinetics, 27 000 cells were seeded at day 0 in a 6-well plate with 2 ml of medium for all conditions. At day 4, medium was changed and at day 7, cells were trypsinized (0.05% trypsin, 0.53mM EDTA; Thermo Fisher Scientific) and this cycle was repeated until D 49. For co-culture, HME_d2GFP cells were mixed 20:80 with HME-RAS_ER_-dsRED cells. For 1 week-kinetics in a 6 well-plate, around 5 000 HME_d2GFP cells were mixed with 22 000 HME-RAS_ER_-dsRED cells per well. Cells were trypsinized at days 7 and 14. For transwell kinetics, 7000 HME_d2GFP cells were plated in the insert for 27 000 HME-RAS_ER_-dsRED cells per well at day 0. Cells were trypsinized at days 7 and 14.

### Mammospheres

Cells were suspended in DMEM-F12 medium and counted. 10 000 cells were seeded into an ultra-low-attachment 24-well plate (Corning). In necessary cases, 4-OHT was added to each well every 4 days (without removing the old medium) and 100 µl of fresh medium was added every 3 to 4 days. Mammospheres were collected after 7–10 days by centrifugation (800g), washed two times in phosphate-buffered saline PBS, trypsinized (100 µl, 30 min) and dissociated mechanically through pipetting. Cells were assessed microscopically for single-cellularity. Next, cells were resuspended with 900 µl of fresh medium and sieved through a 40-μm sieve. 3 rounds of mammospheres formation of 7 to 10 days were conducted. The number of spheres (nonadherent spherical clusters of cells with basal membrane) for each well was evaluated under the microscope after round 3. Quantification of formed mammospheres was performed microscopically by manual counting. Images of mammospheres were acquired using a ZEISS Axio Vert.A1 Inverted Microscope.

### Organoids

Cell suspension was resuspended in Reduced Growth Factor Basement Membrane Matrix BME (BioTechne). A 50 µl drop of this BME suspension containing 1000 cells was placed in the center of a well of a prewarmed ultra-low-attachment 96-well plate (Corning), allowed to harden at 37 °C for 20 minutes. Upon complete gelation, 200 µl of organoid medium supplemented with different factors was added to each well ^59^ The plate was incubated in a humidified 37 °C / 5% CO_2_ incubator. Culture medium was changed every two days. After 10 days, 4-OHT was added to the medium every 3 days in induced RAS condition. After 21 days of 4-OHT treatment or non-treatment, organoids were fixed in buffered formalin for multiplexed immunofluorescence staining.

### Multiplexed immunofluorescence

For histological examination, tissue samples (organoids or cell pellets) were fixed in 10% buffered formalin and embedded in paraffin using the cytoblock kit (Microm Microtech France, E/7401150). 4-µm-thick sections of formalin-fixed, paraffin-embedded tissue were prepared according to conventional procedures.

Multiplexed immunofluorescence was performed on a Bond RX automated immunostainer (Leica biosystems) using OPAL detection kits (AKOYA bioscience). Sequential immunofluorescence stainings were performed using:

-Organoids Panel: OPAL 520 (anti-CK5, Leica #NCL-L-CK5), OPAL 690 (anti-CK18, Agilent #M7010), OPAL 780 (anti-VIMENTINE, Agilent #M0725).
-Cells Panel: OPAL 520 (anti-IL-1 α, LSBio #LS-C33821), OPAL 620 (anti-IL-6, Abclonal #A0286), OPAL 570 (anti-ZEB1, Bethyl #IHC-00419), OPAL 690 (anti-Ki67 Mib1, Agilent #M7240).

Samples were counterstained with DAPI (Sigma Aldrich, D8417). Fluorescent slides were mounted using Prolong™ Gold Antifade Reagent (Invitrogen, Ref# P36930).

Sections were scanned using the Vectra POLARIS device (Akoya bioscience). An autofluorescence treatment of images was carried out using the Inform software (Perkin Elmer). For analysis of cells, an evaluation of the staining was carried out using the HALO Image

Analysis Software (Indica Labs). Cells were considered Ki67-positive if the fluorescence intensity value was > 30, IL-6-positive if the fluorescence intensity value was >25, IL-1α-positive if the fluorescence intensity value was >75 and ZEB1-positive if the fluorescence intensity value was > 32.

### Soft-agar colony formation assay

A 2X DMEM/F12 powder including L-glutamine (Sigma Aldrich #D0547) was re-suspended with sterile water and completed with 1.2 g/L sodium bicarbonate (Sigma Aldrich #S5761). This prewarmed medium was supplemented with 20% heat-inactivated fetal calf serum, 200 UI/mL penicillin, 200 µg/mL streptomycin, 1mg/ml hydrocortisone, 20 ng/ml EGF and mixed 1 :1 with melted SeaPlaque Agarose (Lonza) solution to obtain a final 0.75% low-melting agarose base which was then overlaid with a suspension of cells in 0.45% low-melting agarose (10 000 cells/well). 6-well plates were incubated for 4 weeks at 37°C with 5% CO_2_/95% air, and colonies defined as a group of >20 cells were counted under the microscope. In experiments using GFP-tagged cells, only GFP-positive cells were counted. Experiments were performed in triplicate.

### SA-β-Galactosidase staining

Cells were rinsed with PBS and fixed 4 minutes in 2% formaldehyde/0.2% glutaraldehyde, then washed twice with PBS and incubated at 37°C for 18 hours in a buffer containing 40 mM sodium phosphate (pH 6.0), 5 mM K4Fe(CN)6, 5 mM K3Fe(CN)6, 150 mM NaCl, 2 mM MgCl_2_ and X-Gal re-suspended with N,N-Dimethylformamide at 20 mg/ml and diluted in sterile water to obtain a 1 mg/ml final concentration. All reagents were purchased at Sigma Aldrich except for X-Gal powder (VWR).

### Multiplex cytokine assay

We assessed 31 cytokines in each sample with the Panel (human) kit (Meso-Scale Discovery (MSD)), which measures IL-1α, IL-1β, IL-2, IL-4, IL-5, IL-6, IL-7, IL-8, IL-10, IL-12p40, IL-12p70, IL-13, IL-15, IL-16, IL-17A, GM-CSF, TNF-α, TNF-β, IFN-gamma, TGF-β1, TGF-β2, TGF-β3, IP-10, MCP-1, MCP-4, MIP-1α, MIP-1β, MDC, CCL17, eotaxin 1, eotaxin 3, and VEGF. MSD plates were analyzed on the MS2400 imager (MSD). Calibrator dilutions and samples were prepared according to the manufacturer’s recommendations.

### Reagent, antibodies, and cytokines

To induce RAS activation in HME-derived cells, (Z)-4-Hydroxytamoxifen (4-OHT) (Sigma, #H7904) was added in the medium at 500 nM every 3 to 4 days. Cells were treated with dasatinib (1.5µM) or DMSO every 3 days.

All antibodies except Tocilizumab-Roactemera^TM^ were purchased at R&D Systems (Bio-Techne): IL-1α /IL-1F1 antibody (MAB200), IL-1β /IL-1F2 antibody (MAB201), IL-6 antibody (MAB2061), IL-8/CXCL8 antibody (MAB208), GM-CSF antibody (MAB215) and Normal Human IgG Control (1-001-A). Tocilizumab (Anti-Human IL-6R, Humanized Antibody IgG1; Tocilizumab-Roactemera^TM^ 20 mg/ml, Roche) was stored at 4°C and used at 50 µg/ml final concentration. Other antibodies were reconstituted in aliquots stored at -80°C and once used were stored at 4°C for no more than 2 weeks. Antibodies were used at the following final concentrations guided by MSD/ELISA highest concentrations measured: anti-IL-6 (0.6 µg/ml), anti-IL-8 (1µg/ml), anti-IL-1α (1µg/ml), anti-IL-1β (0.3 µg/ml), anti-GM-CSF (4 µg/ml), and normal human IgG control (50 µg/ml). Medium was changed every 3 to 4 days and antibodies were added at the same time.

Cytokines were purchased at R&D Systems (Bio-Techne): Recombinant Human IL-1α/IL-1F1 Protein (200-LA), Recombinant Human IL-1β/IL-1F2 Protein (201-LB), Recombinant Human IL-6 Protein (206-IL), Recombinant Human IL-8/CXCL8 Protein (208-IL), Recombinant Human GM-CSF Protein (215-GM). These cytokines were reconstituted in sterile PBS with 0.1% bovine serum albumin (Sigma Aldrich) and stored at -80°C. Once thawed, cytokines were kept at 4°C up to 2 weeks. Cytokines were used at the following final concentrations: IL-6 (2 ng/ml), IL-8 (10 ng/ml), IL-1α (1.5 ng/ml), IL-1β (50 pg/ml), GM-CSF (1 ng/ml), normal human IgG control (10 ng/ml). Cytokines were added every 2 days and medium was changed every 3 to 4 days.

### RNA extraction, reverse transcription, and PCR

Adherent cells were washed in PBS (Eurobio scientific) and TRIZOL reagent (Merck Millipore) was added. After sorting, cells were directly centrifuged and washed before TRIZOL was added. Total RNA was isolated using the TRIZOL Reagent according to the manufacturer’s instructions. Total RNA concentration and purity were determined from absorbance at 260 nm and 280 nm using a ND-1000 NanoDrop spectrophotometer. Reverse transcription (RT) was performed with 300 ng to 1 µg of RNA using cDNA RT kit (Thermo Fisher Scientific). Quantitative PCR (qPCR) analysis was performed with 3 µl of a 1:10 dilution of the resulting cDNA. DNA amplification was monitored by real-time PCR with a CFX96 instrument (Bio-Rad) and analyzed with Bio-Rad CFX manager software. The relative quantification of gene expression was performed using the comparative CT method, with normalization of the target gene to HPRT1 housekeeping gene.

List of primer sequences used for Q-PCR analysis:

Human *ZEB1* AGG GCA CAC CAG AAG CCA G and GAG GTA AAG CGT TTA TAG CCT CTA TCA, human *ZEB2* AAG CCA GGG ACA GAT CAG C and GCC ACA CTC TGT GCA TTT GA, human *SNAI1* GCT GCA GGA CTC TAA TCC AGA and ATC TCC GGA GGT GGG ATG, human *SNAI2* TGG TTG CTT CAA GGA CAC AT and GTT GCA GTG AGG GCA AGA A, human *TWIST1* GGC TCA GCT ACG CCT TCT C and CCT TCT CTG GAA ACA ATG ACA TCT, human *TWIST2* CAT GTC CGC CTC CCA CTA and GCA TCA TTC AGA ATC TCC TCC T, *HPRT1* TGA CCT TGA TTT ATT TTG CAT ACC and CGA GCA AGA CGT TCA GTC.

TaqMan qRT–PCR assay was used for detection of mature miRNAs. Reagents, primers and specific probes for miR-200c and miR-141 were obtained from Applied Biosystems. RT reactions and real-time qPCR were performed according to the manufacturer’s protocols from 40 ng of RNA per sample. RNU48 was used as a loading control.

### Immunoblot analysis

Cells were washed in PBS with CaCl_2_ and lysed in a radio-immuno precipitation assay (RIPA) buffer (100 mM NaCl, 1% NP40, 0.1% SDS, 50 mM Tris pH 8) supplemented with protease inhibitor cocktail (tablets, Roche), phenyl-methane sulfonyl fluoride (PMSF, Merck Millipore) and a phosphatase inhibitor cocktail (Merck Millipore). Cells were scratched at 4°C and lysed cells were sonicated. Protein concentration was dosed with Bio-Rad Protein Assay Dye Reagent (BIORAD).

Primary antibodies used include anti-ZEB1 (Merck Millipore # HPA027524), anti-ZEB2 (Merck Millipore #HPA003456), anti-SNAIL+SLUG (Abcam #ab85936), anti-phospho-p44/42 MAPK (Erk1/2) (Thr202/Tyr204) (Cell signalling #4370)), anti-p44/42 MAPK (Erk1/2) (Cell signalling #9102), anti-RAS (G12V mutant specific) (Cell signalling #14412), anti-GAPDH (Merck Millipore #ABS16) and anti-tubulin (Sigma Aldrich #T7816).

Secondary antibodies used include horseradish peroxidase-conjugated goat anti-mouse (Santa Cruz #sc-2005) or mouse anti-rabbit polyclonal antibodies (Santa Cruz #sc-2357). Western blots were revealed using a western blot luminol reagent kit (BIORAD). Luminal signal was detected by ChemiDoc (BIORAD). Post-analysis was done with ImageLab software 6.0.1 (BIORAD).

### Flow cytometry analysis and sorting

For CD44/CD24/CD104 or EpCAM staining, cells were stained with primary antibodies from Miltenyi Biotech: anti-CD44 labelled Fluorescein-5-isothiocyanate (FITC) (clone REA690, dilution 1:400), anti-CD24 labelled phycoerythrin (PE) (clone 32D12, dilution 1:50), anti-CD104 (Integrin β4)-APC (clone: REA236, dilution 1:40) or anti-CD326 (EpCAM)-APC (clone REA764, dilution 1/400). For analyzing GFP- or dsRED-tagged cells, anti CD44-Brilliant Violet 421 (clone BJ18, dilution 1:50; Ozyme) and anti CD24-APC (clone ML5, dilution 1:100; Ozyme) were used.

Fresh cells were washed once in PBS, stained protected from light at 4°C for at least 1 hour and then washed again with PBS 3 times. A FACS Canto II cytometer (Becton Dickinson) or BD LSR Fortessa (Becton Dickinson) was used for data acquisition alongside Diva software. The cytometer was calibrated daily using Cytometer, Set-up and Tracking (CST) beads (Becton Dickinson) according to the manufacturer’s instructions. Sample acquisition was made at a medium flow rate and set to record the highest number and identical number of events possible for each condition. Post-analysis was done using Flowlogic software (Miltenyi Biotec). Antibody validation information is available on the manufacturers’ websites. Cell sorting was performed using BIORAD S3 (Biorad) for GFP- and DSRED-tagged cells after cell infection. For all other sorting experiments, FACS ARIA III (BD Biosciences) was used.

### Statistical analysis

For all biological experiments, significance testing was performed using Prism Software version 7.0 (GraphPad). P values were calculated by one-way analysis of variance (ANOVA) and Tukey multiple comparison test as specified in the figure legends. All experiments were repeated at least 3 times, unless stated otherwise. Kinetics analysis data were plotted as median +/-range and showing all points. P-values <0.05 were considered to be significant. All statistical tests are two-tailed.

### Bioinformatics analysis on breast cancer dataset

#### Samples and statistics

Breast tumour samples used in this study are from METABRIC ^9^ and TCGA Research Network (https://www.cancer.gov/tcga). Cohorts and breast cancer cell lines are from CCLE database ^10^. All analyses, statistical tests and figures were realized using either the R software (version 3.6.1) or GraphPad Prism 8.0 (GraphPad Software Inc., San Diego, USA). All statistical tests are two-tailed.

#### Expression data processing

METABRIC microarray expression data from discovery and validation sets were extracted from the EMBL-EBI archive (EGA, http://www.ebi.ac.uk/ega/; accession number: EGAS00000000083) (Normalized expression data files) ^9^. Normalized expression data per probe of the discovery set and the validation set were combined after independent normalization of each set with a median Z-score calculation for each probe. The expression levels of different probes associated with the same Entrez Gene ID were averaged for each sample in order to obtain a single expression value by gene.

TCGA BRCA RNAseq expression data were extracted as Fragments Per Kilobase Million (FPKM) values from the GDC data portal (https://portal.gdc.cancer.gov/). FPKM data by gene were converted to Transcript Per Million (TPM) as follows: for each gene g ∈ G and each sample s ∈ S,

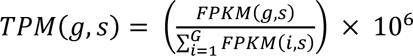

RNAseq expression data from the CCLE breast cell lines were extracted as Reads Per Kilobase Million (RPKM) values from the CCLE data portal (https://portals.broadinstitute.org/ccle). RPKM data by gene were converted to TPM as follows: for each gene g ∈ G and each sample s ∈ S,

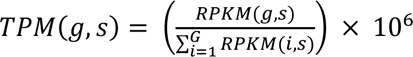

#### Molecular breast cancer subtype assignment

For both tumours and cell lines, attribution of breast cancer molecular subtypes was performed using the R package genefu ^60^. Basal-like, luminal A, luminal B, Her2 and normal-like subtype assignments were computed from 5 different algorithms (PAM50, AIMS, SCMGENE, SSP2006 and SCMOD2) ^61–65^. An assignment was considered final if defined by at least 3 different algorithms. In case of divergence between classifiers, PAM50 subtype attribution was used. For METABRIC and TCGA breast tumours, classification as a claudin-low subtype was defined by nearest centroid method. For this, the Euclidean distance between each tumour sample and the previously described claudin-low and non-claudin-low centroids for tumour samples were determined using the 1,667 genes defined by Prat et al. as significantly differentially expressed between claudin-low tumours and all other molecular subtypes ^5^.

For CCLE breast cancer cell lines, claudin-low status assignment was performed using the nine-cell line predictor via the R package genefu ^60^.

#### Pathway analysis

Single-sample GSEA (ssGSEA) scores were computed through gsva R package ^66^ using gene signatures from the molecular signature database (MSigDB) (msigdbr R package) ^67^. Pan-cancer transcriptomic EMT signature defined by Tan *et al*.^68^ was used to compute EMT scores for each sample.

### Single cell transcriptome analysis

#### Sample preparation

Full-length scRNAseq libraries were prepared using the Smart-seq2 protocol ^69^ with minor modifications. Briefly, freshly harvested single cells were sorted into 96-well plates containing the lysis buffer (0.2% Triton X-100, 1U/µl RNase inhibitor, Thermo Fisher Scientific). RT was performed using SuperScript II (Thermo Fisher Scientific) in the presence of 1 μM oligo-dT30VN (IDT), 1 μM Template-switching oligonucleotides (QIAGEN), and 1 M betaine. cDNA was amplified using the KAPA Hifi Hotstart ReadyMix (Roche) and IS PCR primer (IDT), with 23 cycles of amplification. Following purification with Agencourt Ampure XP beads (Beckmann Coulter), product size distribution and quantity were assessed on a Bioanalyzer using a High Sensitivity DNA Kit (Agilent Technologies). A total of 140 pg of the amplified cDNA was fragmented using Nextera XT (Illumina) and amplified with Nextera XT indexes (Illumina). Products of each well of the 96-well plate were pooled and purified twice with Agencourt Ampure XP beads (Beckmann Coulter). Final libraries were quantified and checked for fragment size distribution using a Bioanalyzer High Sensitivity DNA Kit (Agilent Technologies). Pooled sequencing of Nextera libraries was carried out using a HiSeq4000 (Illumina) to an average sequencing depth of 0.5 million reads per cell. Sequencing was carried out as paired-end reads of 75 bp length (PE75) with library indexes corresponding to cell barcodes.

#### Data processing

For the Smart-seq2 single-cell samples, quality check of the sequenced reads was performed with the FastQC v0.11.8 software to ensure the quality standards. Next, reads were aligned with STAR v2.5.4b on the human genome reference GRCh38 with Gencode 32 annotations. Gene expression was estimated with RSEM v1.3.0

#### Single cell analysis

Fastq files were aligned on the human genome (hg38) using STAR v2.7.0f (default parameters) ^70^. Raw counts for each gene were also computed by STAR, using the gencode V29 annotations.

Counts data for each cell of the same plate were grouped together.

Quality control of the data was done using the R packages scater ^71^ and scran ^72^. To remove low-quality data, we filtered out cells with < 3000 genes or > 7000 genes. We also removed genes expressed in < 5% of cells. No more filtering was needed. Data from all cells was grouped in a single matrix and normalized using the logNormCounts method. Using the 500 most variable genes, we computed PCA and UMAP dimension reductions. We then clustered the data with the buildKNNGraph method (K=15).

Signature scores for each cell were extracted using the ssGSEA method from the GSVA package ^66^ except for the Jean-Paul Thierry EMT signature, for which we used the published method for cell lines ^68^.

We used the FindMarkers method from Seurat ^73^ (using the Model-based Analysis of Single-cell Transcriptomics (MAST) method) to identify gene markers for each data cluster. We then selected the top marker genes from each cluster (p-value < 0.001 and fold change > 1 or < -1) and checked for enriched pathways in these genes using the clusterProfiler package ^74^ and the Hallmark, C2 and C5 genesets from MSigSB ^75^.

We used the Slingshot method ^76^ for trajectory analysis on the previously established data clusters to which PCA was applied for dimension reduction.

## Data availability

The raw FASTQ files and the processed single cell sequencing data of this study can be obtained from Gene Expression Omnibus (GEO). METABRIC data are available in the European Molecular Biology Laboratory-European Bioinformatics Institute (EMBL–EBI) archive (accession number: EGAS00000000083) and from Supplementary Information in ^9^ and ^77^. TCGA BRCA RNAseq expression data were extracted as Fragments Per Kilobase Million (FPKM) values from the GDC data portal (https://portal.gdc.cancer.gov/). RNAseq expression data from the CCLE breast cell lines were extracted as RPKM values from the CCLE data portal (https://portals.broadinstitute.org/ccle).

## Code availability

All original code is publicly available as of the date of the publication.

## Acknowledgements

The authors would like to thank Aruni P. Senaratne for critical reading of the manuscript. We are grateful to Cancer Research Centre of Lyon-Centre Léon Bérard core facilities, the Cell Imaging Platform and the Flow Cytometry Core Facility. We acknowledge the help of Nicolas Gadot and Clementine Leneve (Research Pathology Platform East) for multiplex immunofluorescence stainings, Holger Heyn, Sara Ruiz Gil and Gustavo Rodriguez-Esteban (CNAG-CRG, Barcelona Institute of Science and Technology, Spain) for single-cell analyses. HDB was supported by Fondation pour la Recherche Médicale (FRM) and LabEx DEVweCAN. This work was supported by funding from the Ligue Nationale contre le Cancer (EL2016.LNCC/AIP and EL2019.LNCC/AIP) and SIRIC LYriCAN (INCa-DGOS-Inserm_12563). This work was additionally supported by funding from Inserm within the framework of the International Associated Laboratory between the Cancer Research Centre of Lyon, France, and the Victorian Comprehensive Cancer Centre of Melbourne, Australia (LIA/LEA 2016, ASC17019CSA) and from Le Cancer du sein. Parlons-en! association. The graphical abstract was produced using Servier Medical Art (https://smart.servier.com).

## Author contributions

A.P. and A.P.M. conceived the study, supervised the work and analyzed the data; H.D.B. designed and performed most experiments and analyzed the data; L.T. and R.P. performed the bioinformatics analysis; F.F., C.L., R.B., F.A., B.G. and A.P.M. assisted in experiments; M.O. contributed to discussions about the project and exchanged ideas; A.P wrote the manuscript with inputs from M.O., A.P.M. and H.D.B. All authors discussed the results and commented on the manuscript.

## Corresponding authors

Correspondence to Alain Puisieux or Anne-Pierre Morel.

---

**Extended Data Fig. 1.**
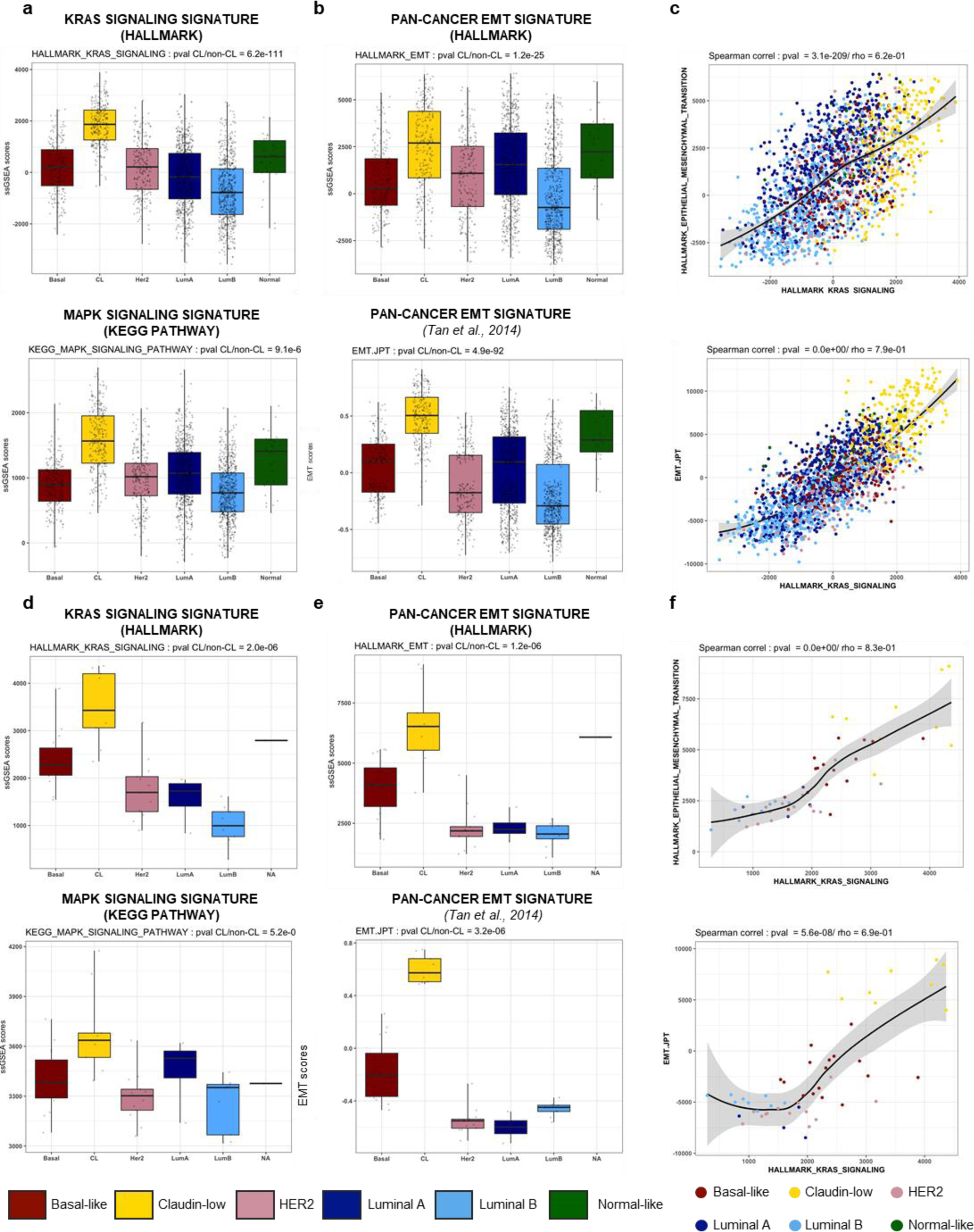
RAS/MAPK and EMT pathways are highly correlated and strongly activated in claudin-low breast cancer subtype. **a**, **b**, Distribution of ssGSEA or EMT scores for each molecular subtype of breast cancer in METABRIC cohort, related to (**a**) RAS or MAPK pathway (ssGSEA score) and (**b**) EMT pathway (ssGSEA or EMT score ^66^); **c**, Spearman correlations between KRAS signalling pathway (ssGSEA score) and EMT pathway (ssGSEA or EMT score ^66^) in breast tumours from the METABRIC dataset; **d**,**e**, Distribution of ssGSEA or EMT scores for each molecular subtype of breast cancer in CCLE cohort, related to (**d**) RAS or MAPK pathway (ssGSEA score) and (**e**) EMT pathway (ssGSEA or EMT score ^66^); **f**, Spearman correlations between KRAS signalling pathway (ssGSEA score) and EMT pathway (ssGSEA or EMT score ^66^) in breast tumours from the CCLE dataset.

**Extended Data Fig. 2.**
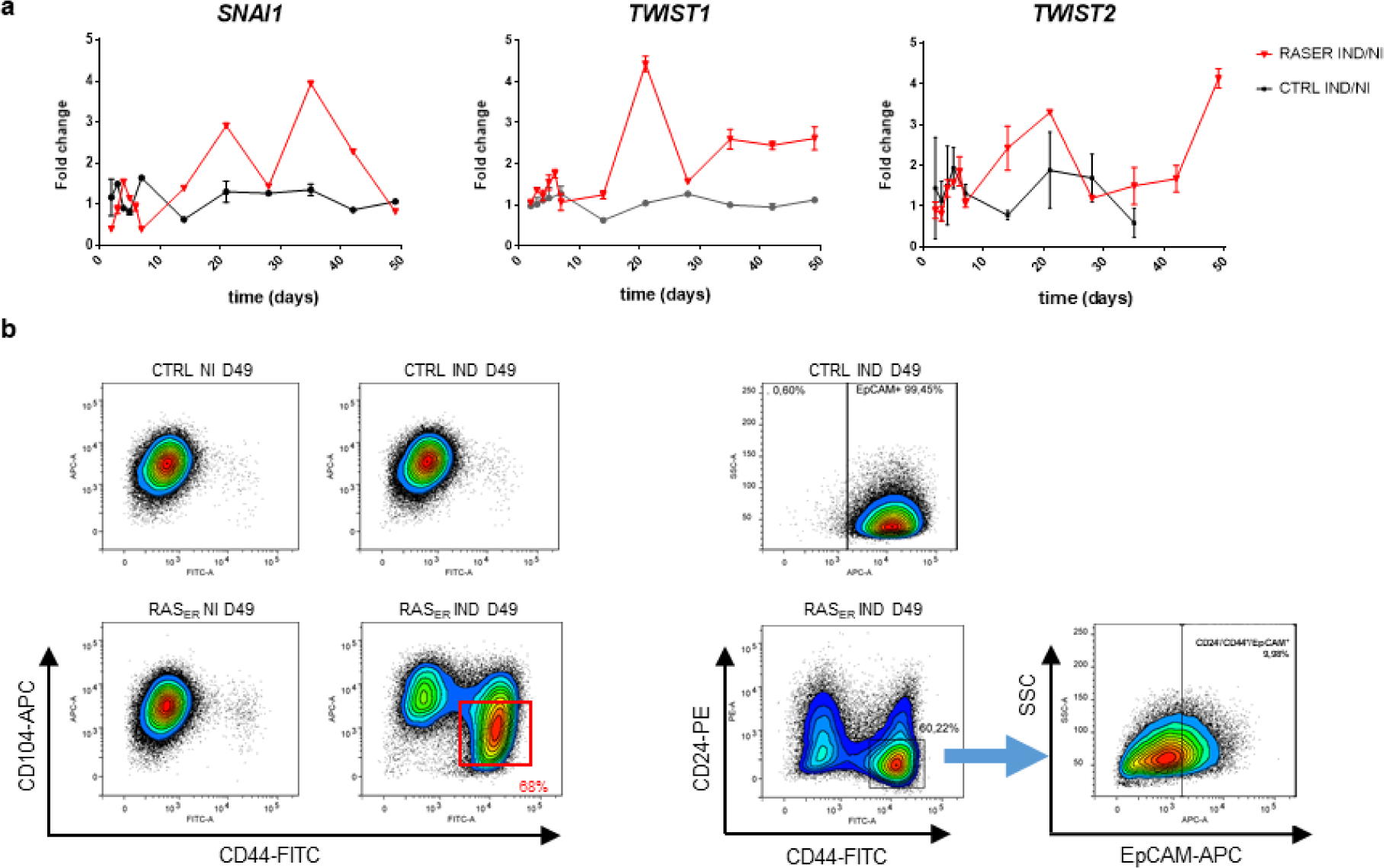
RAS-activation in HME-RAS_ER_ cells induces expression of EMT-TFs and a shift in CD24/CD44/CD104/EpCAM phenotype. **a**, Fold change expression across time of mRNA for EMT-TFs *SNAI1, TWIST1* and *TWIST2* in HME-RAS_ER_ cells or in HME- CTRL cells induced with 4-OHT (IND) or not (NI). Median± range (*n*=2 independent experiments); **b**, Representative flow cytometry analysis of CD24, CD44, CD104 and EpCAM markers in HME-RAS_ER_ (RAS_ER_) and HME-CTRL (CTRL) cells after 49 days of 4-OHT treatment (IND) or not (NI).

**Extended Data Fig. 3.**
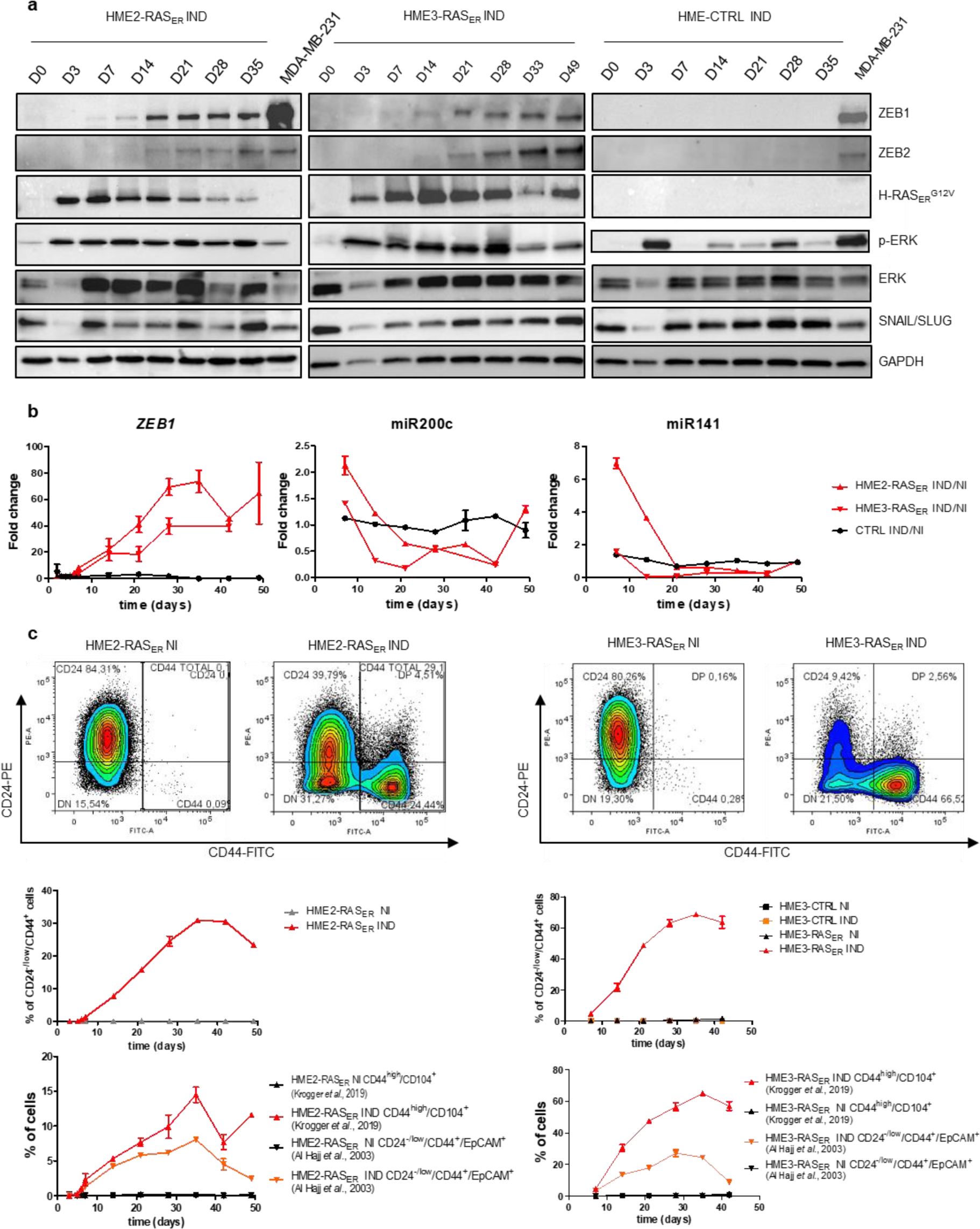
RAS-activation in HME2-RAS_ER_ and HME3-RAS_ER_ cells induces expression of EMT-TFs and a shift in CD24/CD44/CD104/EpCAM phenotype. **a**, Immunoblot showing the expression of HRAS_ER_^G12V^, pERK1/2, ERK, ZEB1, ZEB2 and SNAIL/SLUG in HME2-RAS_ER_ cells, HME3-RAS_ER_ cells and HME-CTRL cells at indicated time points after 4-OHT treatment. MDA-MB 231 cell line was used as positive control for ZEB1, ZEB2 and SNAIL/SLUG expression. GAPDH level was used as loading control; **b**, Fold change expression across time of *ZEB1* mRNA, miR200C and miR141 in HME2-RAS_ER_ and in HME3-RAS_ER_ cells or in HME-CTRL cells induced with 4-OHT (IND) or not (NI). Median± range (*n*=2 independent experiments); **c**, Representative FACS analysis of CD24, CD44, CD104 and EpCAM markers in HME2-RAS_ER_ and HME3-RAS_ER_ cells at day 28 of induction with 4-OHT (IND) or not (NI). Kinetics analysis and quantification of the percentage of CD24^-/low^/CD44^+^ cells, CD24^-/low^/CD44^+^ /EpCAM^+^ cells and CD44^+^/CD104^+^ cells across time after 4-OHT treatment. Median± range (*n*=2 independent experiments).

**Extended Data Fig. 4.**
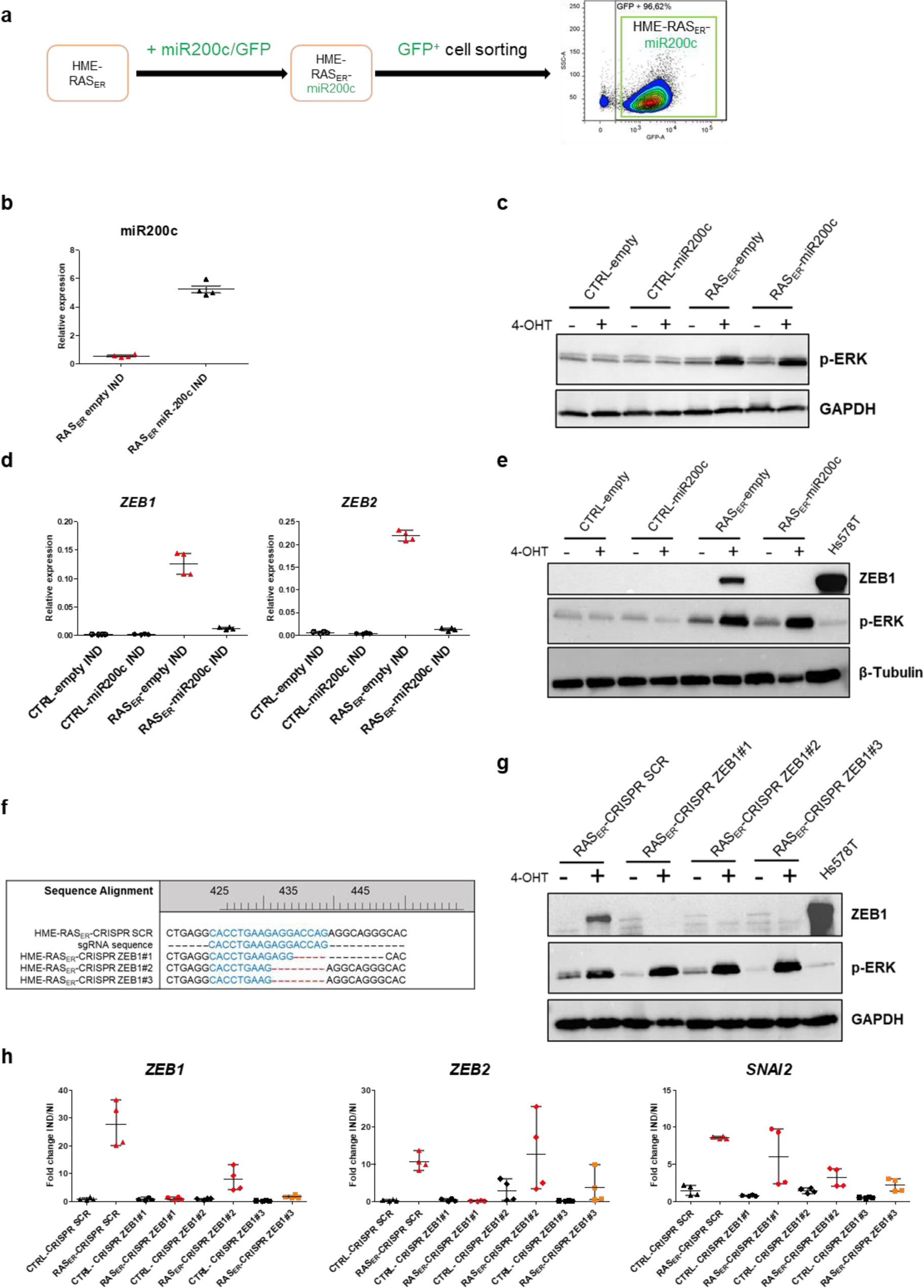
Cellular plasticity and transformation capabilities after RAS activation are ZEB1-dependent. **a**, Experimental outline. miR200c expression was associated with GFP expression. Cell sorting based on GFP^high^ expression in HME-RAS_ER_ miR200c cells and HME-RAS_ER_-empty cells; **b**, miR200c expression in HME-RAS_ER_-miR200c or in HME-RAS_ER_-empty cells induced with 4-OHT (IND). Median± range (*n*=4); **c**, Immunoblotting of pERK in HME-RAS_ER_-miR200c or in HME-RAS_ER_-empty cells treated with 4-OHT (+) or not (-) and analyzed at day 4. GAPDH level was used as loading control; **d**, *ZEB1* and *ZEB2* mRNA expression in HME-RAS_ER_-miR200c or in HME-RAS_ER_-empty cells after 28 days of O-HT treatment (IND). Median±range (*n*=2 independent experiments); **e**, Immunoblotting of pERK and ZEB1 in HME-RAS_ER_-miR200c or in HME-RAS_ER_-empty cells treated with 4-OHT (+) or not (-) and analyzed at day 28. β-Tubulin level was used as a loading control. Hs578T cell line was used as a positive control for ZEB1 expression; **f**, Genomic DNA sequencing by Sanger method for assessing HME-RAS_ER_-CRISPR ZEB1.Sequence alignment of a CRISPR SCR clone and 3 CRISPR ZEB1 KO clones (clones #1, #2 and #3); **g**, Immunoblotting of pERK and ZEB1 in the 3 HME-RAS_ER_-CRISPR ZEB1 clones (#1, #2 and #3) or in HME-RAS_ER_-CRISPR SCR induced with 4-OHT (+) or not (-) and analyzed at day 28. GAPDH level was used as a loading control. Hs578T cell line was used as a positive control for ZEB1 expression; **h**, Fold change in expression (induced/not induced) of *ZEB1*, *ZEB2* and *SNAI2* mRNA in the three HME-RAS_ER_-CRISPR ZEB1 clones (#1, #2 and #3) or in HME-RAS_ER_-CRISPR SCR induced with 4-OHT or not for 28 days. Relative expression was determined by the ΔΔCt method, normalized to *HPRT1* expression and divided by the expression of the untreated sample at the same time point. Median±range (n=2 independent experiments).

**Extended Data Fig. 5.**
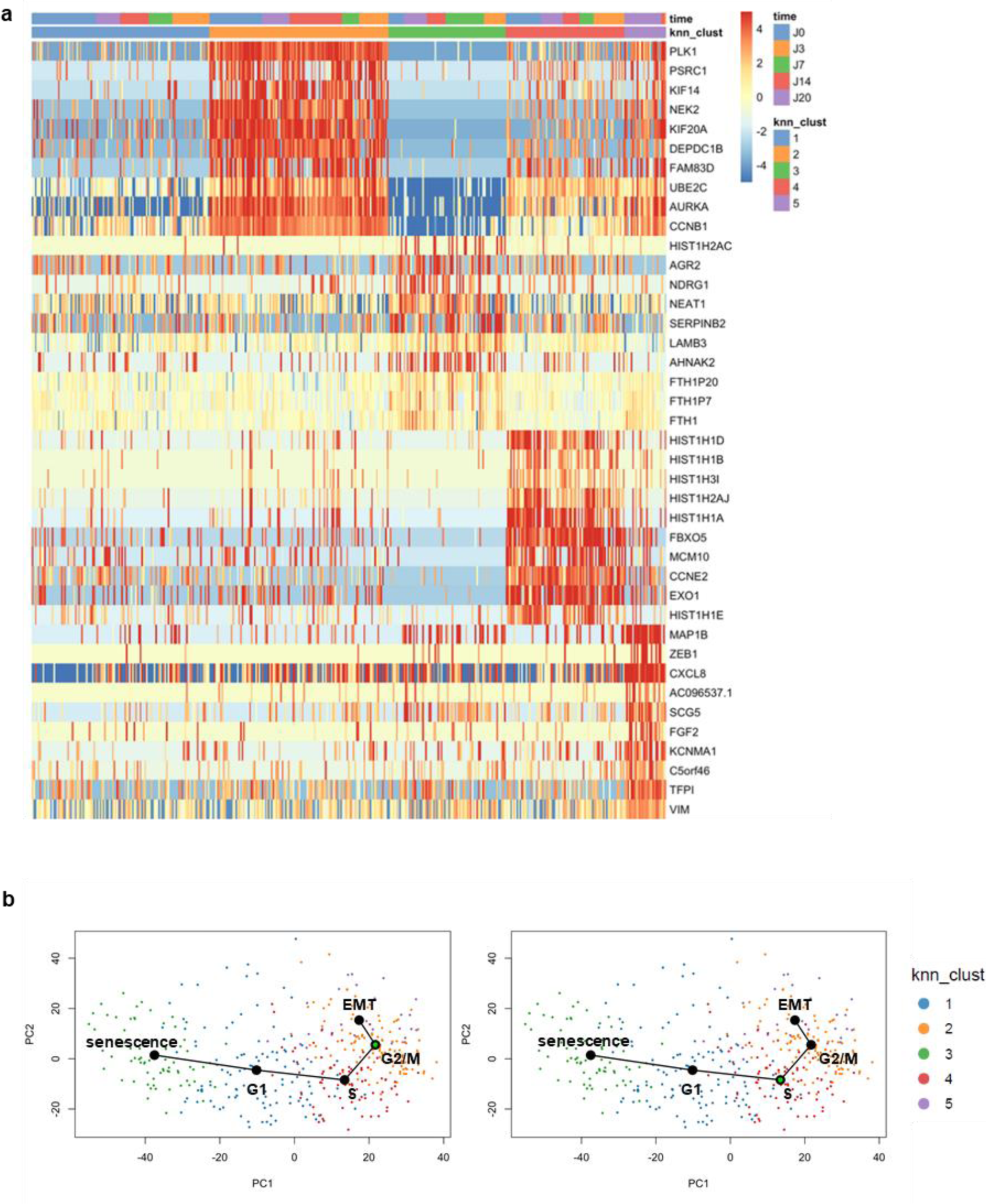
Identification of distinct senescence and EMT clusters in induced HME-RAS_ER_. **a**, Heatmap representing the expression of the 10 most up-regulated genes of each cluster compared to all other clusters; **b**, Projection of the trajectory analysis performed by the Slingshot algorithm along the first two axes of an unsupervised PCA of all cells. Cells are colored by clusters. Upper panel: Starting point of the trajectory analysis is cluster 2 (green circle), Bottom panel: Starting point of the trajectory analysis is cluster 4 (green circle).

**Extended Data Fig. 6.**
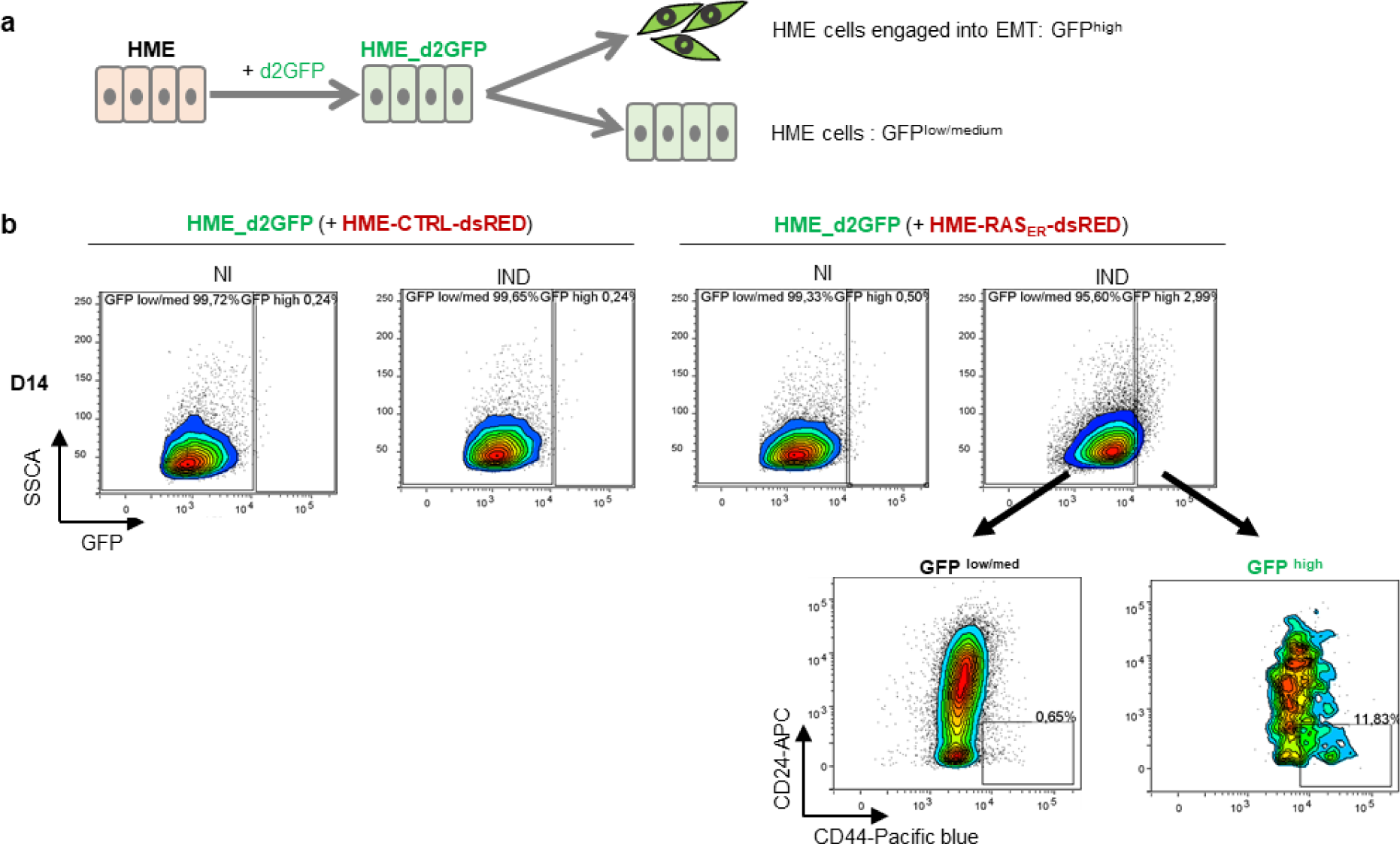
ZEB1-dependent cellular plasticity is driven by a paracrine mechanism. **a**, Experimental outline of HME_d2GFP generation and expected read out; **b**, Representative FACS analysis of CD24^-/low^/CD44^+^ population according to GFP intensity (GFP ^high^ or GFP ^low/med^) of HME-d2GFP cells co-cultured with HME-RAS_ER_-dsRED cells or HME- CTRL-dsRED cells induced with 4-OHT (IND) or not (NI) for 14 days.

**Extended Data Fig. 7.**
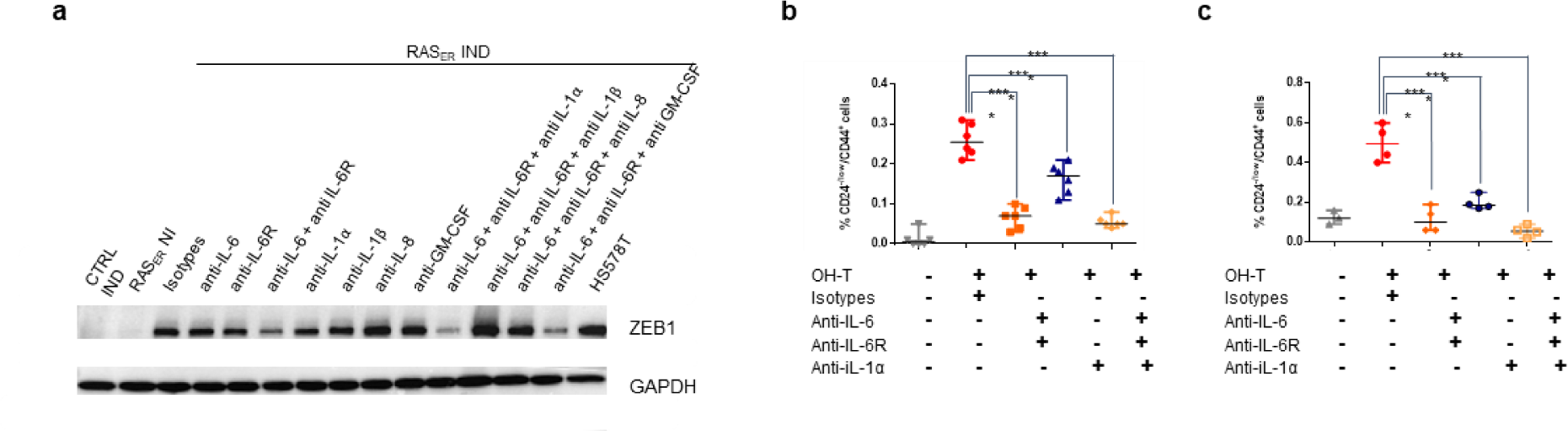
ZEB1-dependent plasticity is driven by cytokines IL-6 and IL-1α secreted from senescent cells. **a**, Immunoblotting of ZEB1 expression in HME-CTRL or HME-RAS_ER_ cells (RAS_ER_) induced by 4-OHT (IND) or not (NI), treated with neutralizing antibodies or isotype controls and analyzed at day 28. Data is representative of three independent experiments (*n*=3); **b**, Quantification of CD24^-/low^/CD44^+^ population from HME- d2GFP cells co-cultured with HME-RAS_ER_-dsRED cells induced with 4-OHT (+) or not (-), treated with indicated neutralizing antibodies or isotype controls and analyzed at day 20. Median±range (*n*=7); **c**, Quantification of CD24^-/low^/CD44+ population from HME-d2GFP cells cultured in transwell plate with HME-RAS_ER_-dsRED cells induced with 4-OHT (+) or not (-), treated with indicated neutralizing antibodies or isotype controls and analyzed at day 20. Median±range (*n*=4).

